# Nuclear speckle proteins form intrinsic and *MALAT1*-dependent microphases

**DOI:** 10.1101/2025.02.26.640430

**Authors:** Min Kyung Shinn, Dylan T. Tomares, Vicky Liu, Avnika Pant, Yuanxin Qiu, You Jin Song, Yuna Ayala, Gregory W. Strout, Kiersten M. Ruff, Matthew D. Lew, Kannanganattu V. Prasanth, Rohit V. Pappu

## Abstract

Nuclear speckles are enriched in serine / arginine rich splicing factors (SRSFs), such as SRSF1. Splicing factors and proteins such as TDP-43 concentrate into distinct speckle territories to enable pre-mRNA processing. We have discovered that SRSFs and TDP-43 are block copolymers and the protein-specific interplay of inter-block repulsions and attractions drives spontaneous microphase separation. This gives rise to size-limited, ordered assemblies, that are 30 – 45 nm in diameter. Depending on the protein, each microphase comprises several tens to hundreds of molecules. The sub-micron scale territories observed in cells are shown to be clusters of microphases. The regulatory lncRNA *MALAT1* binds preferentially to SRSF1 microphases to enhance microphase separation and alter microphase structures. Microphase separation enables the concentration of finite numbers of splicing factors into assemblies with distinct nanoscale structures that can be modulated by *MALAT1*. Our findings provide a structural framework for the functional organization of splicing factors.

## Introduction

Nuclear speckles are ubiquitous in metazoan cells ^1^. They function as hubs that regulate key steps of gene expression, including transcription, pre-mRNA processing, and RNA export from the nucleus^1–8^. Speckles are enriched in spliceosomal RNA binding proteins that contribute to pre-mRNA splicing ^8–13^. The activities of splicing factors within speckles are regulated by polyadenylated RNAs and regulatory long non-coding RNAs (lncRNAs) that are enriched in the transcriptomes of speckles^7,14–16^. At this juncture, the driving forces that help organize splicing factors into distinct territories within speckles remain unclear.

It has been proposed that nuclear speckles form via liquid-liquid phase separation (LLPS) ^17^. Although the proteins SON and SRRM2, which are localized to the cores of speckles, form condensates with signatures of LLPS ^18,19^, the overall structural organization of speckles raises questions about the validity of LLPS as the organizing process. For example, super resolution structured illumination microscopy (SIM) has shown that most speckle-associated proteins and lncRNAs are organized in an inhomogeneous manner around a SON-SRRM2 core ^20^. Furthermore, distinct components appear to accumulate into distinct, demixed, sub-micron scale territories ^18,20–25^. Comparison of SIM images to other physical systems suggests that the overall organization within speckles is most likely that of microphases organized around a core that forms via macrophase separation ^26^. To clarify these distinctions, we first define macrophase versus microphase separation and distinguish them from one another.

LLPS is an example of macrophase separation. Macromolecules that drive macrophase separation typically feature uniformly attractive interactions that can be modeled as effective homopolymers ^27^ (**Figure 1A**). The attractive interactions result from the overall incompatibility with the solvent ^28,29^ (**Figure 1A**). The micron-scale sizes of macrophases are orders of magnitude larger than the nanoscale dimensions of the underlying macromolecules. At equilibrium, macrophases are likely to encompass millions of molecules and grow via coarsening to form a single dense phase that coexists with the dilute phase (**Figure 1A**) ^30^. Coarsening is seldom observed in cells, and this has been explained as the result of active processes ^31,32^, solid particles such as Pickering agents ^33^, elastic networks ^34,35^, or dynamical arrest ^36^ that suppress coarsening.

**Figure 1.**
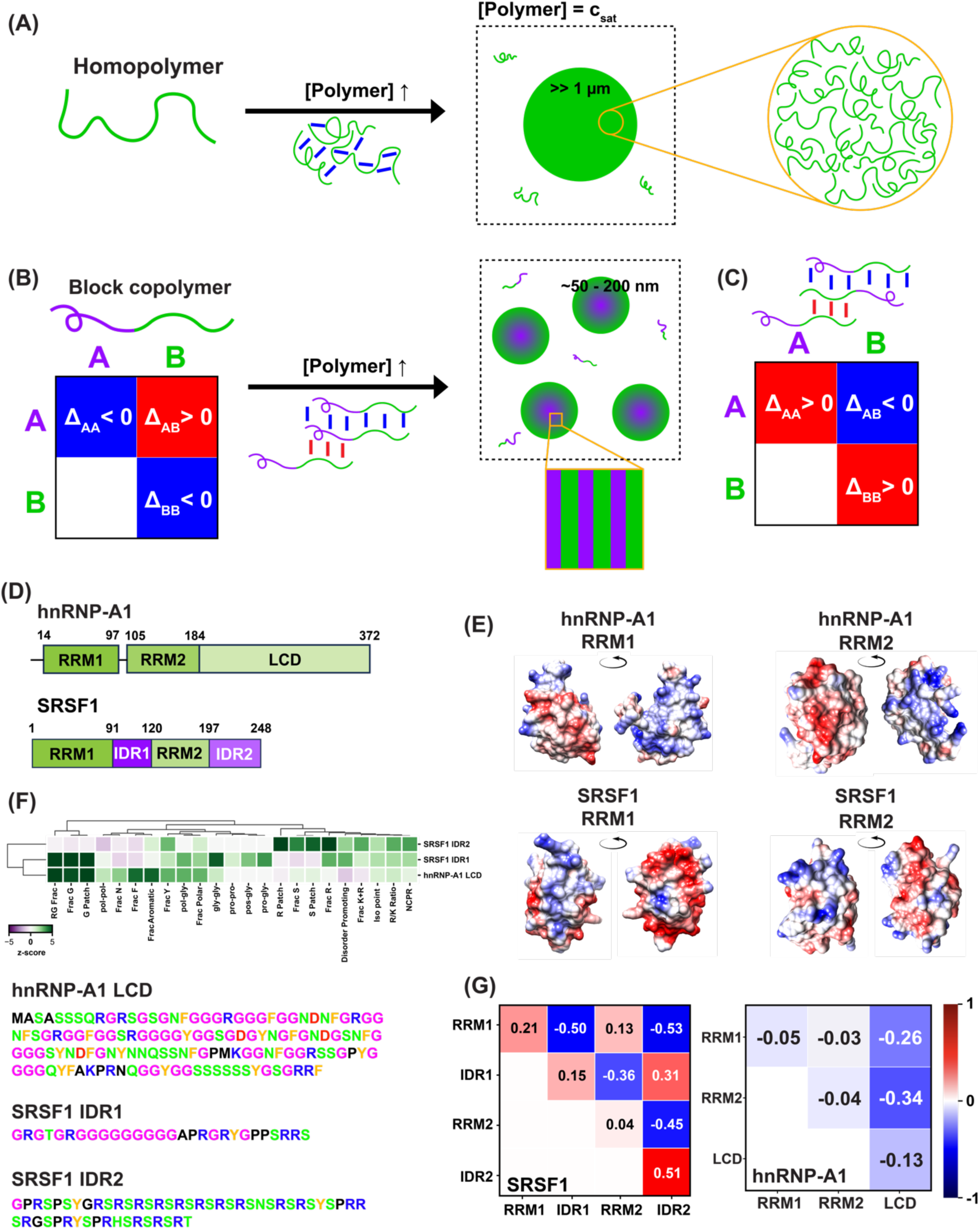
SRSF1 is a block copolymer and a candidate for microphase separation. **(A)** Schematic of a homopolymer that undergoes macrophase separation. Homopolymers or effective homopolymers ^27^ are defined by uniformity of interactions (blue lines) that are on the order of thermal energy. Above a threshold concentration of the homopolymer, known as the saturation concentration (c_sat_), a single micron-scale dense phase forms that coexists with the dilute phase. The inset shows a schematic of how some of the millions of molecules might be organized within the dense phase. **(B)** Schematic of a di-block copolymer featuring A-(purple) and B-blocks (green). A normalized interaction coefficient Δ_XY_ is used to define the interactions between blocks. In the interaction matrix, the homotypic inter-block interactions Δ_AA_ and Δ_BB_ are attractive, whereas the incompatibility of the blocks leads to repulsive heterotypic interactions denoted by positive values of Δ_AB_. **(C)** An alternative scenario for the di-block system is one where the homotypic interactions are repulsive, positive values of Δ_AA_ and Δ_BB_, whereas heterotypic interactions are attractive (negative values of Δ_AB_). **(D)** Sequence architectures of hnRNP-A1 and SRSF1 showing the domain boundaries. **(E)** Mapping of the computed surface electrostatic potentials (SEPs) onto the surfaces of the RRMs from hnRNP-A1 and SRSF1. Regions in blue are positive potentials, regions in red are negative potentials, and regions in white are neutral. The magnitudes and signs of the site-specific SEPs are shown in **Figure S1**. **(F)** Non-random sequence features within the LCD of hnRNP-A1 and IDR1 and IDR2 of SRSF1. The color-coded sequences from which the features were extracted are shown below the NARDINI+ analysis. The colors correspond to polar residues (green), positively charged residues (blue), negatively charged residues (red), glycine residues (magenta), aromatic residues (orange), and others (black). The inventory of sequence features is provided in the **Methods** section. A positive z-score implies either an enrichment of a compositional bias when compared to the human proteome or a non-random linear segregation of specific pairs of amino acid types. A negative z-score implies either a depletion of a compositional bias when compared to the human proteome or a non-random linear mixing of specific pairs of amino acid types. **(G)** Interaction coefficients Δ_XY_ computed from atomistic simulations for each pair of distinct domains within SRSF1 (left) and hnRNP-A1 (right).

A different explanation for the formation of supramolecular assemblies that do not coarsen comes from connections to a process known as microphase separation ^37–42^. This process is driven by macromolecules with block copolymeric architectures ^39,41,43,44^ (**Figure 1B**) where distinct blocks feature distinct chemistries ^41^. The simplest of such systems are di-block copolymers. Here, the homotypic (between the same types of blocks) inter-block interactions are strongly attractive, and the heterotypic (between different types of blocks) inter-block interactions are strongly repulsive ^39,43–46^ (**Figure 1B**). Inter-block interaction patterns defined by homotypic repulsions and heterotypic attractions (**Figure 1C**) can also lead to microphase separation. At finite concentrations, the inter-block attractions can drive phase separation, but these interactions must compete against repulsions. The competition between inter-block attractions and repulsions results in assemblies known as microphases that are of fixed size featuring specific nanoscale morphologies and internal ordering ^39,44^. Microphase sizes, structures, internal ordering, and the number of molecules per microphase are determined by the number of distinct blocks ^41^, the sizes of distinct blocks, and the inter-block interaction patterns of block copolymers (**Figure 1B, 1C**) ^47^.

Here, we report that the speckle-associated SR family splicing factors SRSF1, SRSF3, SRSF5, SRSF7, and TDP-43 (transactive response DNA binding protein) undergo intrinsic and RNA-dependent microphase separation. The RRMs (RNA recognition motifs) and intrinsically disordered regions (IDRs) of the SRSFs and TDP-43 represent distinct types of domains and sequence blocks. Each of these proteins form microphases that are 30-45 nm in diameter depending on the protein or mixture of proteins. The regulatory lncRNA *MALAT1* (Metastasis-Associated Lung Adenocarcinoma Transcript 1) is enriched at the periphery of speckles ^20,48,49^ and harbors distinct binding motifs for SRSF1 ^49–51^. We report that *MALAT1* modulates the driving forces for microphase separation and the structures of microphases formed by RRM-containing, speckle-enriched SRSF1 and speckle-associated TDP-43 ^50^.

## Results

### Establishing the block copolymeric features of SRSF1

The protein hnRNP-A1 is a known driver of macrophase separation ^52,53^. It is a nuclear protein that localizes to cytoplasmic stress granules during stress ^53^. Like SRSF1, hnRNP-A1 comprises two RRMs, and the overall architectures are similar between the two proteins (**Figure 1D**). However, the four RRMs across the two proteins are quite different from one another. The overall pairwise sequence identities are low (less than 35% as determined by multiple sequence alignment ^54,55^ and their atomic, site-specific surface electrostatic potentials (SEPs) are different (**Figure 1E**, **S1A, S1B**). For example, the median SEP is +3.82 kcal/mol**·***e* for RRM2 of SRSF1, while it is-0.11 kcal/mol**·***e* for the RRM2 of hnRNP-A1. In contrast, the SEP is +0.30 kcal/mol**·***e* for RRM1 of SRSF1, and +0.94 kcal/mol**·***e* for RRM1 of hnRNP-A1. This implies that the surfaces of the RRMs are differently amphoteric across the different proteins. We also computed the hydrophobicity of each of the RRMs according to their sequence (**Figure S1C, S1D**). Hydrophilic linear sequence regions are defined by negative values, and hydrophobic regions are defined by positive values. On average, the two RRMs of hnRNP-A1 have more hydrophobic regions than the RRMs of SRSF1. These calculations suggest that despite similarities in the overall folds, the surface-mediated interactions are likely to be very different for each of the four RRMs.

In addition to the RRMs, both proteins feature IDRs. The C-terminal low complexity domain (LCD) of hnRNP-A1 is the site of several pathogenic mutations ^53,56^. On its own, the A1-LCD undergoes macrophase separation ^27,57,58^. SRSF1 features two IDRs: IDR1 is a 28-residue Gly-rich region between the two RRMs, whereas IDR2, located at the C-terminal end, is a 53-residue-long region enriched in Ser/Arg residues (**Figure 1F**). We used the NARDINI+ algorithm ^59,60^ to compare the sequence features within the three IDRs. These comparisons were quantified in terms of z-scores, which were obtained by comparing the sequence features within each IDR to the human IDRome ^60,61^. IDR1 in SRSF1 and the LCD of hnRNP-A1 are enriched in Gly-rich patches, Gly content, and the presence of RG-motifs (**Figure 1F**). On the other hand, the two regions differ in the Phe, Asn, and aromatic contents (enriched in A1-LCD), and the extent of linear segregation of Gly residues (high in IDR1 of SRSF1). Furthermore, the IDR2 of SRSF1 has a distinct sequence grammar when compared to the IDR1 of SRSF1 and the LCD of hnRNP-A1 ^62^. The standout sequence feature is the presence of Ser/Arg repeats, which is quantified as enrichments of Ser/Arg residues (Frac S, Frac R, R Patch, S Patch).

The question is if the differences in RRMs and IDRs across the two proteins translate into quantifiable differences in inter-domain interaction patterns. To answer this question, we used atomistic simulations (see **Methods**) to characterize the strengths and types of homotypic and heterotypic inter-domain interactions for the two proteins. The domains could be a pair of RRMs, a pair of IDRs, or an RRM and an IDR. For each pair of domains labeled X and Y, we extracted an inter-domain interaction parameter designated as Δ_XY_. This parameter is normalized and bounded such that-1 ≤ Δ_XY_ ≤ +1. Attractive inter-domain interactions are characterized by negative values of Δ_XY_ whereas positive values reflect repulsive interactions. The magnitudes of Δ_XY_ quantify the strengths of the attractions / repulsions. If |Δ_XY_| ≈ 0, then the inter-domain attractions and repulsions are counterbalanced, implying that X and Y are ideal, non-interacting domains.

Results from simulations of inter-domain interactions of hnRNP-A1 and SRSF1 show the following: For hnRNP-A1, all the Δ_XY_ values are either negative or near zero (**Figure 1G**). Therefore, hnRNP-A1 lacks any inter-domain repulsions, and the strongest attractions are mediated by the A1-LCD. The lack of inter-domain repulsions that can compete with the attractions implies that hnRNP-A1 behaves more like a homopolymer that undergoes macrophase separation ^27,52,53^. Conversely, for SRSF1 the homotypic interactions are either strongly repulsive (RRM1-RRM1, IDR1-IDR1, IDR2-IDR2) or nearly ideal (RRM2-RRM2) (**Figure 1G**). The heterotypic IDR-RRM interactions (IDR1-RRM1, IDR1-RRM2, IDR2-RRM1, and IDR2-RRM2) are attractive, whereas the heterotypic IDR1-IDR2, and RRM1-RRM2 interactions are repulsive. These interaction patterns suggest that SRSF1, unlike hnRNP-A1 is a complex tetra-block copolymer ^41,63^ featuring different patterns of inter-block (in)compatibilities. Next, we characterized the phase behavior of SRSF1 in vitro to assess whether it forms microphases or macrophases.

### SRSF1 has a measurable threshold concentration for forming large assemblies

We measured right-angle light scattering (RALS) in vitro to quantify the existence of a threshold concentration for microphase separation (c_µ_) of SRSF1. In the absence of a change in assembly state, the scattering intensity should increase linearly with the protein concentration ^64^. The presence of a discontinuity in the light scattering intensity, measured over a range of protein concentrations, indicates the formation of a distinct phase defined by large assembly sizes. We measured scattering intensities of SRSF1 over a 100-fold concentration range and observed a discontinuity at 0.45 ± 0.05 µM SRSF1 in an aqueous buffer (10% glycerol, 20 mM HEPES, pH 7.4, 50 mM KCl, 5 mM MgCl_2_) (**Figure 2A**) ^50^.

**Figure 2.**
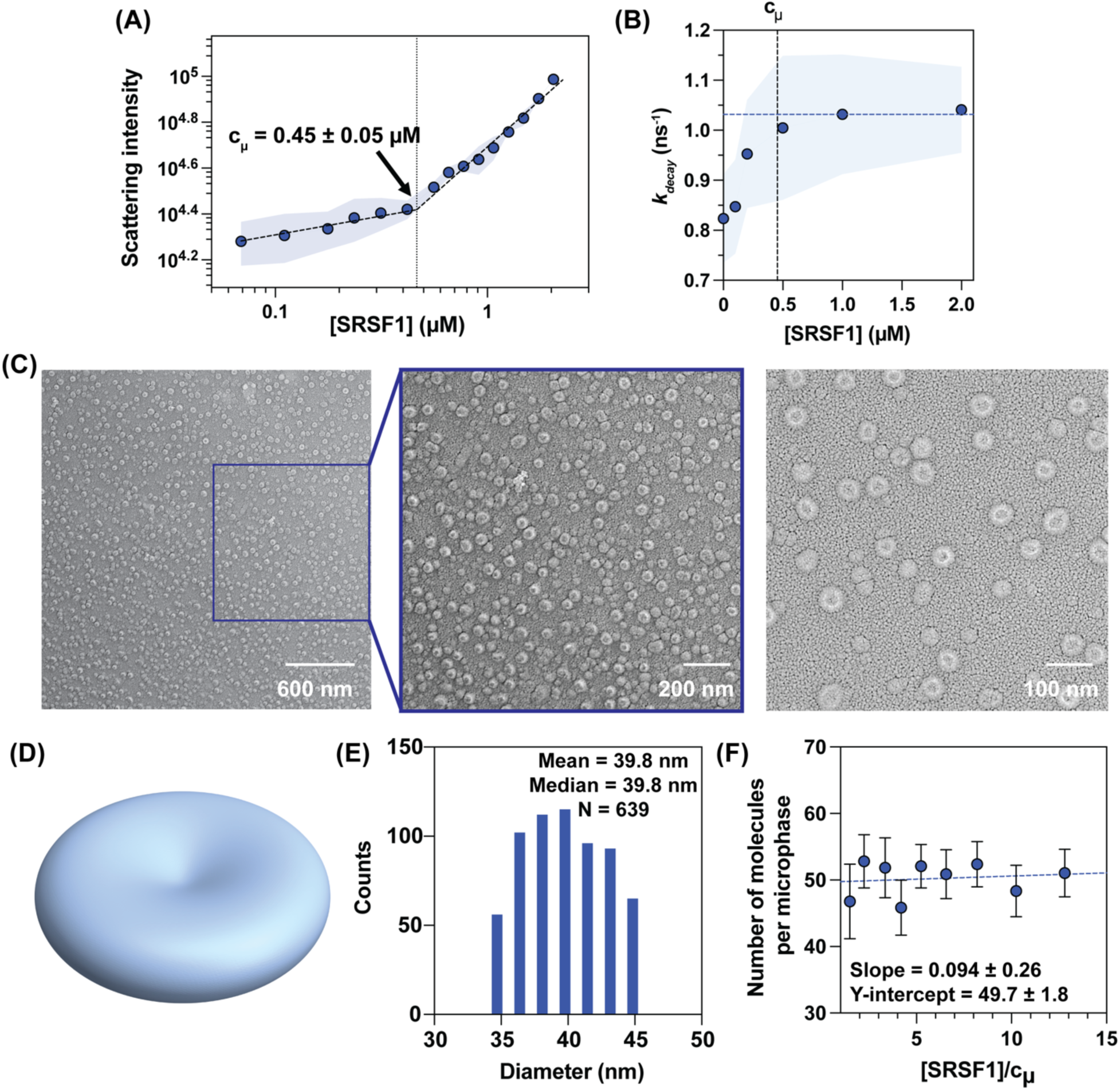
SRSF1 forms microphases of distinct biconcave structures. **(A)** Data from RALS measurements for SRSF1. The scattering intensity is plotted as a function of SRSF1 concentration. Results are shown from a single representative measurement (solid circles joined by dashed lines) and the spread over three technical replicates across different protein preps (pale blue envelope). The discontinuity in the scattering intensities at 0.45 ± 0.05 µM is denoted as c_µ_. **(B)** Decay rate constants (*k*_decay_) of the fluorescence lifetime of pyrene. The solid circles are rate constants from a single representative measurement. The pale blue envelope shows the range of values obtained from three technical replicates. The rate constant shows a plateauing behavior (horizontal dashed line) near the c_µ_ (∼0.45 µM) estimated from RALS. The increase in the rate constant beyond the plateau value is proportional to the increase in the concentration of microphases with increasing [SRSF1]. **(C)** QF-DEEM images at three different magnifications show biconcave morphologies of SRSF1 microphases and their reproducibility across a large field-of-view. **(D)** Three-dimensional rendering of the morphologies observed in panel **(C)**. The method of rendering, described in **Methods**, uses a diameter of 8 units, the biconcave (dent) thickness as 0.5 units, and the maximum thickness of the biconcave structure near the rim as 2 units. **(E)** Histogram of diameters (in nm) constructed from the analysis of 639 distinct images. **(F)** Results from TRFQ experiments were fit as described in the **Methods**. The intercept along the ordinate is 49.7 ± 1.8 molecules, and it quantifies the number of SRSF1 molecules within a microphase at the c_µ_.

The value of c_µ_ was found to be 0.56 ± 0.09 µM in the presence of 150 mM KCl and 5 mM MgCl_2_. The measured c_µ_ increased with increasing concentration of KCl (**Figure S2A**). The parsimonious interpretation is that the threshold concentration beyond which detectably large assemblies form in solution is driven, in part, by complementary electrostatic interactions that are most likely heterotypic inter-domain interactions (**Figure 1G**). Here, we focused mainly on sequence-intrinsic interactions. Accordingly, all subsequent measurements, for all the systems we studied, were performed in solutions comprising 5 mM MgCl_2_ and 50 mM KCl (10% glycerol, 20 mM HEPES, pH 7.4).

We used measurements of pyrene lifetimes as a function of SRSF1 concentration as an independent probe of the presence of a threshold concentration. Microphases can be viewed as block copolymeric analogs of micelle-forming systems ^65^. For the latter, one can measure a critical micelle concentration by monitoring changes in the fluorescence lifetime of pyrene, which is sensitive to the presence of micelles ^65^. We measured fluorescence intensities and lifetimes of pyrene at a fixed pyrene concentration of 20 mM and increasing concentrations of SRSF1 (**Figure S2B**). The pyrene lifetimes, quantified as a single rate constant *k*_decay_, extracted from the decay of pyrene fluorescence as a function of SRSF1 concentrations, increased with SRSF1 concentration and reached a plateau value of 1.04 ± 0.08 ns^-1^ near the c_µ_ (**Figure 2B**). Taken together, two different methods showed the existence of a threshold concentration c_µ_ for SRSF1, and the estimates of c_µ_ using the two methods were similar to one another.

### Structural characterization reveals that SRSF1 forms microphases

Microphases are likely to be ordered on the nanoscale and detection of this type of order requires cryo-electron microscopy or related methods that afford the requisite spatial resolution. However, given the surfactant-like nature of block copolymers, the sample preparation must not create artifacts due to air-water interfaces ^39^. The optimal method turned out to be quick-freeze deep-etch electron microscopy (QF-DEEM) wherein by pressing the sample against a polished copper block cooled by liquid He at 4K ^66^. This approach helped avoid drying, wicking, or exposure to the air-water interface during sample preparation. The flash frozen sample was fractured at-104C/170K to optimize etching, deposition, and imaging of the species of interest ^67^. Recently, Fargason et al. ^68^ reported that SRSF1 is insoluble in a buffer containing 50 mM KCl in the absence of glycerol. They by measured absorbance at 280 nm following ammonium sulfate precipitation and centrifugation. Our measured c_µ_ of 0.45 µM is below the limit of detection by absorbance.

Using QF-DEEM imaging, we observed that above the measured c_µ_, SRSF1 molecules assembled into biconcave-disc-like nanoscale structures (**Figure 2C, 2D**). These structures are reminiscent of red blood cells. We analyzed QF-DEEM images spanning a large field-of-view to quantify the diameters of 639 SRSF1 microphases (**Figure S2C**, see **Methods**). The calculated mean and median values for the diameters of the nanoscale assemblies were identical (39.8 nm) (**Figure 2E**). From atomistic simulations, the computed radius of gyration and mean end-to-end distance of SRSF1, accounting for the conformational heterogeneity of disordered regions, are 2.7 ± 0.2 nm (**Figure S2D**) and 5.3 ± 1.8 nm (**Figure S2E**), respectively. The formation of nanoscale structures above c_µ_ that are an order of magnitude larger than the intrinsic size of an SRSF1 molecule, shows that SRSF1 undergoes microphase separation above a threshold concentration of 0.45 µM ^43^.

Next, we estimated the number of SRSF1 molecules per microphase. While the resolution in QF-DEEM images is high enough to quantify the sizes of the microphases, it is insufficient for resolving the internal organization of the molecules. Therefore, we estimated the numbers of SRSF1 molecules within microphases using time-resolved fluorescence quenching (TRFQ) experiments with pyrene as a fluorescent probe in the presence of cetylpyridinium chloride as a quencher ^69^. The concentration of microphases was estimated from the measured fluorescence decay over a range of SRSF1 concentrations. A 20-fold molar excess of quencher to pyrene ensured that pyrene excimers could not form ^69^. This combined with knowledge of c_µ_ allowed us to estimate the number of SRSF1 molecules per microphase (see **Methods**). Analysis of the TRFQ method yielded an estimate of ∼50 SRSF1 molecules per microphase. Furthermore, we observed a weak dependence of the number of molecules per microphase on the protein concentration with a slope near zero (slope = 0.094) (**Figure 2F**). This implies that the number of molecules remains almost constant over a 13-fold increase in SRSF1 concentration above the c_µ_.

Overall, the data reported thus far show the presence of a threshold concentration c_µ_, which combined with the QF-DEEM data show that SRSF1 assembles into nanoscale structures that are ∼40 nm in diameter. The sizes of the nanoscale structures are only an order of magnitude larger than the dimensions of individual molecules, and these structures are concordant with being microphases, each encompassing ∼50 SRSF1 molecules.

Overall, unlike macrophases, microphases are defined by a narrow distribution of sizes (tens of nm), encompassing tens-to-hundreds, as opposed to millions, of molecules per microphase. Based on the structural parameters, we estimated the effective concentration of SRSF1 molecules within microphases using the formula c_eff_ = (N/V)×(1/N_A_). Here, N =50 is the number of SRSF1 molecules within each microphase, V = 3.35×10^-23^ m^3^ is the volume of each microphase assuming we set the radius to be 2×10^-8^ m, and N_A_ is Avogadro’s number. This calculation yields an estimate of ∼2.47 M for c_eff_ within each microphase. The implication is that each microphase is highly concentrated in a finite number of SRSF1 molecules. The estimate for c_eff_ contrasts with the lower, millimolar concentrations ^27,58^ and millions to tens of millions of molecules that make up micron-scale macrophases ^28,29,70^.

### SRSF1 microphases can associate to form sub-micron scale clusters

Each SRSF1 molecule carries a significant excess of positive charge (+20*e*), with cationic residues making up ∼21% of the sequence. Nanoscale colloidal particles such as microphases, that carry a net charge, can associate to form clusters of fixed size via short-range attractions and long-range repulsions known as SALR ^71–73^. In the case of SRSF1 microphases, counterions in the solution are likely to accumulate around microphases to reduce the buildup of surface charge that derives from the charge-rich, polyelectrolytic IDRs (**Figure 3A**). As protein concentrations increase, the abundance of microphases will also increase. This will enable short-range attractions that drive clustering of microphases, whereby condensed counterion layers are shared among microphases within a cluster ^74^ (**Figure 3A**). The buildup of charge within clusters of microphases engenders long-range repulsions, thereby stabilizing the sizes of clusters. Clustering of microphases via SALR interactions can be destabilized by the addition of a suitable co-ion, ideally one that is identical to cationic residues. Addition of the co-ion should lead to the dispersal of clusters of microphases while maintaining the microphases. We tested these hypotheses using a combination of RALS, direct measurements of cluster sizes in solution, confocal microscopy, and single-molecule imaging.

**Figure 3.**
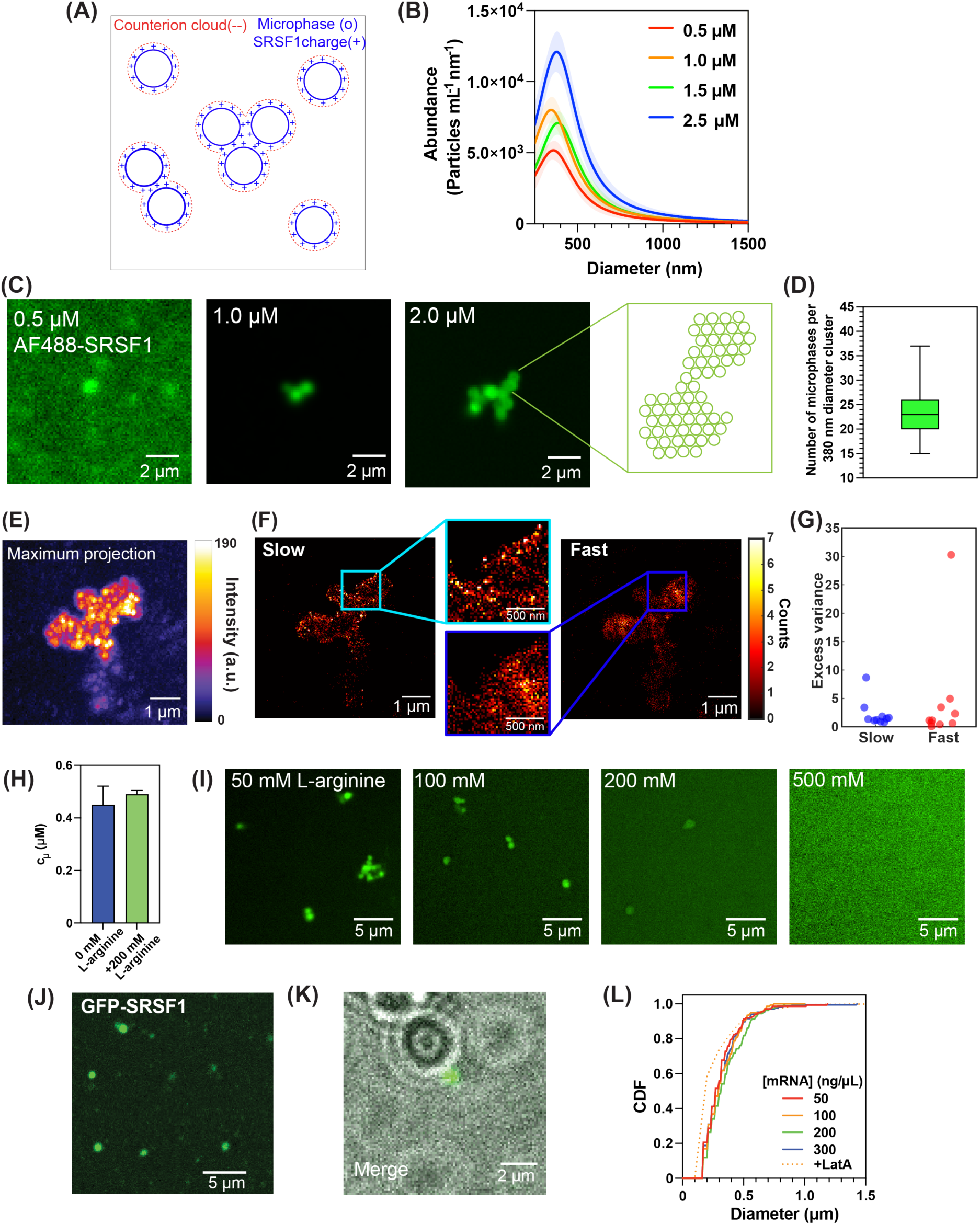
SRSF1 microphases associate to form sub-micron scale clusters. **(A)** Schematic showing how clustering of SRSF1 microphases might arise via SALR-like interactions. An accumulation of surface charge shown by blue + marks will draw a counterion cloud (dashed red envelope) around each microphase (solid blue circles). Increasing the concentration of microphases increases the likelihood that microphases cluster via effective short-range attractions that result from the sharing of counterion clouds among pairs and higher-order clusters of microphases. The attractions saturate beyond a certain size because the buildup of charge creates long-range repulsions. Hence, clusters of microphases do not grow beyond a specific size. **(B)** MRPS measurements of SRSF1 above the c_µ_ are shown as average and standard deviation about the mean. The data show peaks at a constant position while the amplitude increases with increasing concentration of SRSF1. The implication is that the mean sizes stay the same, while the abundance of the clusters increases with increasing concentrations of SRSF1. Results are shown from a single representative measurement (solid lines) and the spread over three technical replicates (pale envelopes). **(C)** Confocal microscopy at concentrations above c_µ_. With increased SRSF1 concentration, the abundance of SRSF1 clusters increases. The schematic depicts how microphases form clusters by touching to form incipient networks of clusters. **(D)** Box and whisker plot from analysis of QF-DEEM images yielded estimates for the number of microphases within a ∼380 nm sized cluster. The box extends from the 25^th^ to 75^th^ percentiles. The line in the middle of the box is at the median value of 23 microphases per cluster. The whiskers extend from the minimum to the maximum number of microphases detected in the probe spheres. **(E)** Single-molecule microscopy with photoactivatable JaneliaFluor 549 (PA-JF549)-labeled SRSF1 shows that the clusters are composed of multiple microphases. The image is produced as a maximum projection of 200 frames collected where the color bar shows the fluorescence intensity. **(F)** Histograms of fluorophores with trajectories of less than 50 nm (slow) or more than 50 nm (fast) over each 20 ms frame for 20,000 frames each with an inset for a zoomed-in field of view of the outlined area. The histograms are represented as heatmaps mapped onto the image. **(G)** Excess variance (see **Methods**) computed over bins that quantify the displacements of slow-versus fast-moving molecules. **(H)** RALS extracted c_µ_ values in the absence and presence of 200 mM L-arginine in experimental buffer are within the error. **(I)** Confocal microscopy images of AF488-SRSF1 in the presence of 50 mM, 100 mM, 200 mM, and 500 mM L-arginine in the experimental buffer. **(J)** GFP-tagged SRSF1 is expressed in *X. laevis* GVs by microinjection of mRNA and SRSF1-positive bodies are observed. **(K)** Merge of images collected simultaneously via brightfield microscopy and the intensity in the GFP channel (see **S3J**) shows that the SRSF1 bodies are juxta identifiable Cajal bodies in the GVs. **(L)** Cumulative distribution functions (CDF) of the diameters of SRSF1-positive bodies remain essentially invariant at different levels of the injected mRNA. Latrunculin A (Lat-A) treatments (orange dotted) to oocytes injected with 100 ng/µL (orange) mRNA did not result in an increase in the size of the intra-GV SRSF1-positive bodies.

Clusters of microphases that form via SALR should have a fixed size, and each cluster should comprise a fixed number of microphases. We measured size distributions in solutions at SRSF1 concentrations above c_µ_ using microfluidic resistive pulse sensing (MRPS). This technique measures the change in electrical resistance as particles of different size pass through a constriction of defined resistance ^75^. The abundance and the size of particles can be determined based on the transit times and the magnitudes of the electrical signals, respectively. Over a 5-fold range in SRSF1 concentrations, all of which were above c_µ_, the MRPS measurements revealed the presence of clusters within a size range of ∼360 nm in diameter. The cluster sizes varied between 359 ± 20 nm and 386 ± 30 nm depending on the concentration. Within error, these values were essentially the same. While the sizes of clusters stayed essentially the same, their abundance increased as the SRSF1 concentration increased (**Figure 3B**).

The sizes of clusters of SRSF1 microphases, as gleaned from MRPS, were above the diffraction limit. Therefore, we used confocal microscopy and SRSF1 molecules that were labeled with AlexaFluor 488 at the N-terminus for direct imaging of the clusters (**Figure 3C, S3A**). We observed clusters of SRSF1 molecules that were ∼400 nm in diameter, which is equivalent to the sizes measured using MRPS. The clusters appeared at SRSF1 concentrations that were equal to or greater than c_µ_. Consistent with MRPS measurements, the sizes of clusters did not change as SRSF1 concentrations were increased. Instead, within every field-of-view (**Figure 3C, S3A**), the abundance of clusters increased with increasing SRSF1 concentration. The clusters appeared to touch without coalescing, forming incipient networks. This too is in accord with expectations at higher concentrations of clusters that form via SALR ^73^.

To arrive at an estimate of the number of microphases per cluster, we used the measured cluster sizes from MRPS measurements as a guide and analyzed distinct fields-of-view in QF-DEEM images. Specifically, we counted the number of microphases within a probe circle of 380 nm in diameter that was tiled across a large field-of-view (**Figure S3B**). Using this analysis, we estimated a median number of ∼23 microphases within a cluster that is of diameter ∼380 nm (**Figure 3D**).

Next, we examined the dynamics of the microphases within clusters using fluorescence recovery after photobleaching (FRAP) of AlexaFluor 488-labeled SRSF1. We observed that the fluorescence intensity did not recover for at least 120 seconds after photobleaching (**Figure S3C**). The small sizes of microphases necessitated a different approach from the diffraction-limited probe of molecular dynamics such as FRAP. So, we utilized single-molecule localization microscopy ^76^ where we measured the blinking of fluorescently labeled SRSF1 molecules.

SRSF1 molecules were labeled with the photoactivatable Janelia Fluor 549 (PA-JF549) at the N-terminus and mixed with unlabeled SRSF1 at a 1:50 molar ratio. This choice allowed for each detected fluorophore to correspond to a single microphase since we determined that a SRSF1 microphase comprises ∼50 molecules (**Figure 2F**). In the single-molecule measurements, we collected 2×10^4^ frames per field-of-view at 20 ms per frame for a total of 6.7 minutes. The maximum projection of all frames onto a single image showed the presence of clusters, which mirrored what was observed with confocal imaging (**Figure 3E, 3C**). Each structure, corresponding to a cluster of size ∼400nm diameter, also contained multiple fluorophores (**Figure 3E**). This was consistent with our analysis of the QF-DEEM images that ∼20-30 microphases are present per cluster.

Next, we utilized single-particle tracking photoactivated localization microscopy (sptPALM) ^77^ to track the motions of SRSF1 molecules within microphases and clusters to measure their displacements in frames that were 20 ms in duration. The tracking was performed by generating a mask for each cluster (**Figure S3D**). Within each mask, the pixels corresponding to single molecules were sorted based on their displacements over 20 ms time intervals. The cumulative distribution function (CDF) of displacements, computed over all fields of view, showed that the probability of finding molecules with displacements that are less than or equal to 50 nm is 0.5 (**Figure S3E**).

The CDF allowed for the classification of molecules as being slow moving (**Movie S1**) if the displacement was confined to being less than 50 nm within a 20 ms frame. Note that this length scale is equivalent to the size of a single SRSF1 microphase, as observed from the line profiles across a region containing the fluorophores (left panel in **Figure S3F**). Molecules were defined to be fast moving if the displacements were larger than 50 nm (**Movie S2**). This likely involves motions between microphases over the 20 ms timescale. Comparing the histograms for the two sets of displacements, we observed that there were more slow molecules, which implies that SRSF1 molecules were mostly confined within microphases (**Figure 3F**).

Next, we computed the excess variance of displacements across all bins within a mask. This was quantified as the variance of displacements within a bin divided by the number of bins within a mask minus one (see **Methods**). The excess variance should be close to zero if the variance of displacements is identical across all bins. However, we observed prominent outliers with some bins showing a strong localization of fluorophores within a ∼50 nm region (**Figure 3G, S3G**). The length scale of this strong localization, and the numbers of fluorophores within the length scale are consistent with the detection of individual microphases.

Finally, we asked if the clusters of microphases could be disrupted by changes to solution conditions, specifically the addition of a co-ion such as L-arginine ^78–80^. Addition of 200 mM L-arginine, while maintaining the pH of the buffer at 7.4, did not alter the driving forces for microphase separation, as assessed by an identical value for the measured c_µ_, using RALS, to in the absence of L-arginine (**Figure 3H, S3H**). However, the addition of an increasing concentration of L-arginine eliminated clustering as observed by confocal microscopy (**Figure 3I**). Fargason et al. ^68^ observed an increase in solubility in the presence of an octameric peptide of RS repeats (RSRSRSRS) or a mixture of arginine and glutamate (Arg/Glu) in solution. In this study, solubility was measured by absorbance at 280 nm following ammonium sulfate precipitation and centrifugation. However, microphases could have remained in the supernatant if clustering of microphases had been disrupted in the presence of the RS peptide or the Arg/Glu solution, as it has been reported that “nanoclusters” of 100-200 nm in diameter need to be subjected to ultracentrifugation to be separated from the solution ^81^.

The effect of adding L-arginine can be rationalized as follows: The counterion cloud around the surfaces of microphases, driven by a buildup of SR-rich IDRs, drives the clustering of microphases through the sharing of counterion clouds ^74^. SALR-like interactions, with long-range repulsions derived from a buildup of charge, will stabilize the sizes of clusters. Addition of L-arginine as a solute competes for the counterions that are shared among microphases which destabilizes clustering and disperses the microphases. However, microphase separation, which is driven mainly by the pattern of inter-block interactions, is not affected by the addition of L-arginine. Overall, the in vitro studies show that SRSF1 forms microphases and that these can associate to form size-limited, sub-micron scale clusters. We tested this prediction in live cells using the *X. laevis* system.

### SRSF1 forms size-limited clusters in cells

We leveraged the large sizes of the *X. laevis* oocytes and their germinal vesicles (GVs) to quantify the sizes of assemblies that SRSF1 forms in GVs. For this, we varied the copy numbers of mRNAs of GFP-tagged SRSF1 by microinjection into oocytes (**Figure S3I**). Following expression, the GVs were extracted and imaged using confocal microscopy ^82^. We observed more than 200 SRSF1-positive bodies in each GV (**Figure 3J**). Some of the SRSF1-positive bodies were attached to a larger non-fluorescent body as observed by brightfield microscopy (**Figure 3K, S3J**). This is consistent with nuclear speckles being attached to Cajal bodies ^83^. The sizes of SRSF1-positive bodies were found to be narrowly distributed with a mean diameter that was less than 500 nm (**Figure 3L, S3K**). This was true irrespective of the injected levels of mRNA, which varies the expression levels of SRSF1 in GVs. The sizes of SRSF1-positive bodies observed in GVs are consistent with the sizes of clusters of SRSF1 microphases observed in vitro.

Nuclear actin can stabilize the sizes of nuclear bodies that form via macrophase separation. These include nucleoli and histone locus bodies (HLBs) ^82^. Feric and Brangwynne showed that nucleoli and HLBs fuse and coarsen upon incubation of *X. laevis* GVs with Latrunculin A (Lat-A), which is a drug that disrupts actin ^82^. However, unlike nucleoli and HLBs, the size distributions of SRSF1-positive bodies did not change when the GVs were treated with Lat-A (**Figure 3L**, orange dashed curve). Therefore, the uniform size distributions observed for SRSF1-positive bodies in oocyte GVs, which are consistent with MRPS data in vitro, appear to be intrinsic properties of the clusters of SRSF1 microphases. They are not the result of stabilization by the nuclear actin network. Taken together with the in vitro, the in-cell data suggest that the sub-micron scale SRSF1 bodies that form in cells are clusters of microphases.

### SRSF3, SRSF5, and SRSF7 are distinct types of block copolymers

Next, we investigated the phase behaviors of other SRSFs. There are twelve human SRSFs named SRSFx where x ranges from 1-12. We chose three additional SRSFs – SRSF3, SRSF5, and SRSF7 – as representative SRSFs with different molecular architecture. In HeLa cells, SRSF1 is the most abundant of the SRSFs followed by SRSF3, and SRSF7 ^84^. SRSF5 is of relatively low abundance, although its architecture shares similarities with SRSF1 (**Figure S4A**). Like SRSF1, SRSF5 has two RRMs. The median SEP of RRM1 of SRSF5 is lower (-2.14 kcal/mol**·***e*) than the RRM2 (0.33 kcal/mol**·***e)* (**Figure S4B**). The median values and distributions of hydrophobicity of the RRMs also differed between the two proteins (**Figure S1C**, **S4C**). Unlike SRSF1 and SRSF5, SRSF3 and SRSF7 have only one RRM each (**Figure S4A**). These RRMs have median SEP values above zero, sharing similarities with the RRM1 from SRSF1. The median hydrophobicity values below zero were similar to the other RRMs examined (**Figure S4C**).

The key differences among SRSF1, SRSF3, SRSF5, and SRSF7 lie in the IDRs (**Figure 4A**). We utilized NARDINI+ to extract the enriched sequence features within the SRSF IDRs. All C-terminal IDRs were found to be enriched in arginine and serine features compared to the human IDRome (red box in **Figure 4A**). However, the lengths of these IDRs vary, with SRSF7 having the longest SR-rich IDR and SRSF1 having the shortest.

**Figure 4.**
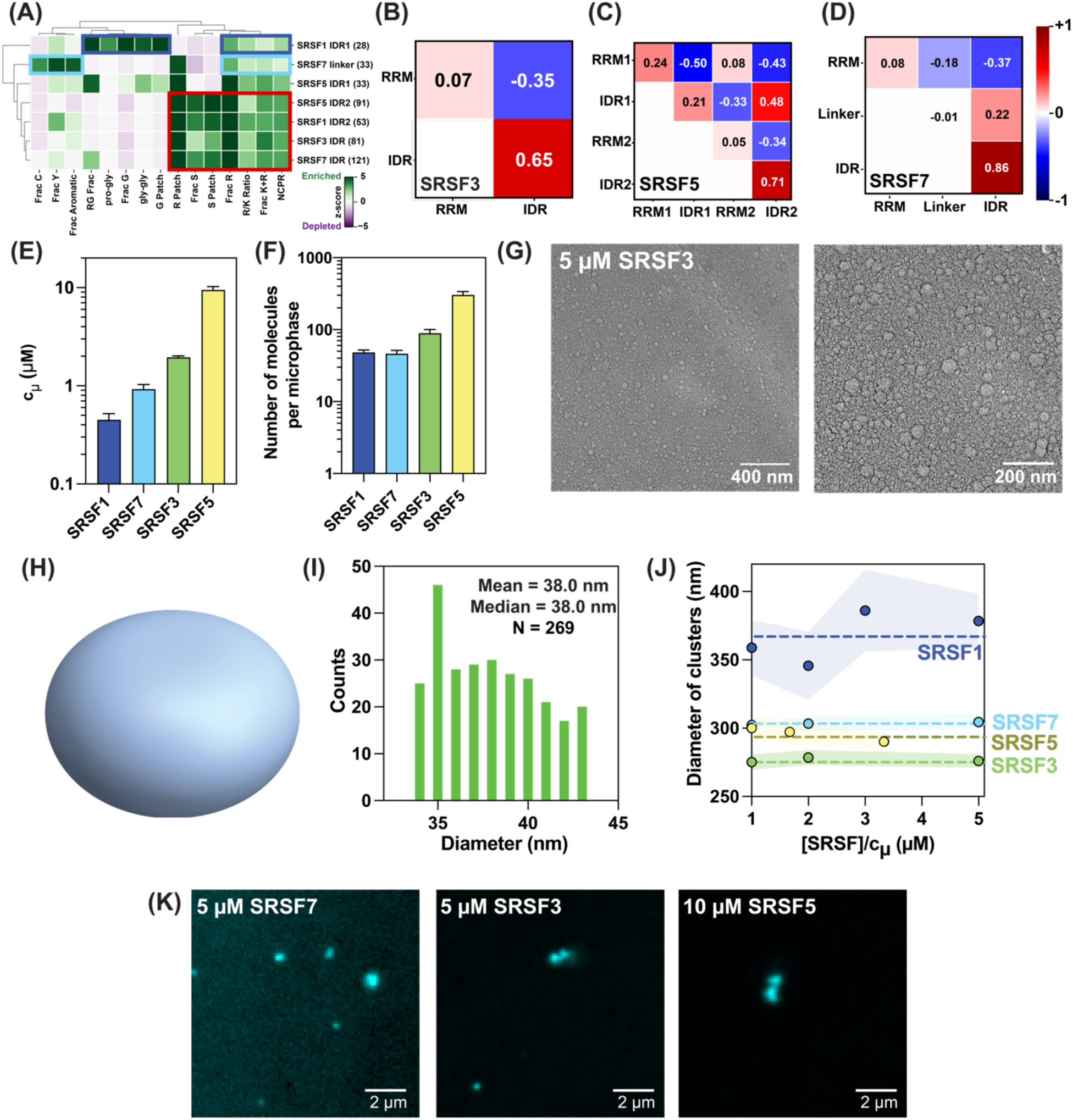
Microphase separation is a shared feature of SRSFs. **(A)** Results from NARDINI+ analysis comparing the sequence features of IDRs in SRSF1, SRSF3, SRSF5, and SRSF7. For clarity, only the sequence features in which at least one sequence had a z-score greater than or equal to 3 are shown. The Ser / Arg rich IDRs (C-terminal IDR in SRSF3 and SRSF7 and IDR2 in SRSF1 and SRSF5) share similarities in terms of compositional biases toward Ser and Arg and the presence of Ser patches (red box). The two distinct IDRs are IDR1 of SRSF1 and the linker between the N-terminal RRM and C-terminal IDR of SRSF7. **(B)** Inter-domain interaction matrix of Δ_XY_ coefficients for SRSF3. **(C)** Inter-domain interaction matrix of Δ_XY_ coefficients for SRSF5. **(D)** Inter-domain interaction matrix of Δ_XY_ coefficients for SRSF7. **(E)** Bar plot of threshold concentrations c_µ_ for SRSF3, SRSF5, and SRSF7 extracted from RALS measurements. The data for SRSF1 are from the RALS measurements shown in Figure 2 and these are incorporated into the bar plot for comparison. The values of c_µ_ increase in the order of SRSF1 (0.45 ± 0.05 µM) < SRSF7 (1.0 ± 0.14 µM) < SRSF3 (2.0 ± 0.14 µM) < SRSF5 (9.5 ± 0.71 µM). **(F)** The number of molecules per microphases determined from TRFQ experiments for the four SRSFs (see **Figure S4F** for values). **(G)** QF-DEEM images of 5 µM, SRSF3 which is above the measured c_µ_, shows a spheroidal morphology. **(H)** Three-dimensional rendering of the morphologies observed in panel **(G)**. The method of rendering, described in the **Methods**, uses a diameter of 8 units, the biconcave (dent) thickness as 5 units, and the maximum thickness of the biconcave structure near the rim as 5 units. **(I)** Histogram of diameters (in nm) constructed from the analysis of images of 269 distinct particles. **(J)** Summary of results from MRPS measurements of SRSFs over a range of protein concentrations. The abscissa is normalized by the intrinsic c_µ_ of each protein for comparison across the SRSFs. The ordinate shows the diameter of clusters in nanometers. The average diameter is 355 nm for SRSF1 (dark blue), 295 nm for SRSF7 (light blue), 298 nm for SRSF5 (yellow), and 276 nm for SRSF3 (green). **G.** Sub-micron-scale clusters of microphases are observed by confocal microscopy for SRSF7, SRSF3, and SRSF5.

Thus, the arginine valence and net charge vary between C-terminal IDRs, which can modulate both homotypic and heterotypic inter-domain interactions. Additionally, SRSF1 IDR1 and the 38-residue linker between the RRM and C-terminal IDR of SRSF7 have enriched sequence grammars that are shared with prion-like domains ^27^. Specifically, SRSF1 IDR contains a glycine tract and the SRSF7 linker is enriched in aromatic residues, including tyrosine.

Next, we used atomistic simulations to characterize the inter-domain interactions for each of SRSF3, SRSF5, and SRSF7 and assessed if they fit the description of being block copolymers (**Figure 4B-4D**). The homotypic inter-domain interactions in SRSF3 are nearly ideal (RRM-RRM) or repulsive (IDR-IDR) whereas the IDR-RRM interactions are attractive (**Figure 4B**). The inter-domain interaction patterns for SRSF5 are qualitatively identical to those of SRSF1. However, the homotypic inter-IDR2 repulsions are stronger in SRSF5 (**Figure 4C**) when compared to SRSF1 (**Figure 1G**). Although the RRM-RRM, RRM-IDR, and IDR-IDR interactions for SRSF7 are qualitatively equivalent to those of SRSF3, the presence of the linker introduces a new degree of freedom into SRSF7 when compared to SRSF3 (**Figure 4D**). Based on the interaction patterns, it follows that SRSF5, like SRSF1, is a tetra-block system, whereas SRSF3 and SRSF7 are di-and tri-block systems, respectively. This should give rise to protein-specific driving forces for microphase separation, and structures that are protein-specific. We tested this, by characterizing the phase behaviors of SRSF3, SRSF5, and SRSF7.

### SRSF3, SRSF5, and SRSF7 undergo protein-specific microphase separation

We performed RALS measurements to quantify the SRSF-specific threshold concentrations (c_µ_). The measured values were as follows: 1.0 ± 0.14 µM for SRSF7, 2.0 ± 0.14 µM for SRSF3, and 9.5 ± 0.71 µM for SRSF5 (**Figure 4E, S4D**). The fluorescence lifetimes of pyrene were also measured over a range of concentrations for each of the three SRSFs. At the c_µ_ for each of the SRSFs, the rate constants plateaued, albeit to protein-specific values of 1.01 ± 0.1 ns^-1^, 0.93 ± 0.17 ns^-1^, and 1.08 ± 0.17 ns^-1^ for SRSF7, SRSF3, and SRSF5, respectively (**Figure S4E**).

Using the TRFQ assay, we estimated there to be ∼46, ∼89, and ∼308 molecules in the SRSF7, SRSF3, and SRSF5 microphases, respectively (**Figure 4F** and **S4F**). The number of molecules did not increase significantly for SRSF7 and SRSF3 with slopes of 2.2 and 4.2 molecules per micromolar increase in protein concentration, respectively. For SRSF5 we obtained a slope of 20.8 molecules per micromolar increase in protein concentration, indicating microphase growth over 9-fold increase in protein concentration. A slope that is smaller than the error in the estimate of the intercept is concordant with microphases increasing in abundance with increased concentration but the number of molecules per microphase being invariant. The slopes for each of the three systems satisfy this criterion.

Next, we used QF-DEEM to probe the structures of microphases formed by SRSF3 as a representative di-block system that could be compared to the microphases formed by the tetra-block SRSF1. QF-DEEM analysis showed that SRSF3 formed spherical as opposed to biconcave structures (**Figure 4G, 4H**). The mean and median diameters were computed to be 38 nm, and these sizes are equivalent to that of SRSF1 (**Figure 4I**). However, the overall morphology is different, and the number of molecules within SRSF3 microphases is double the number of molecules in SRSF1 microphases. This points to different packing patterns within the microphases formed by the di-block system.

Using MRPS (**Figure 4J, S4G**) and confocal microscopy (**Figure 4K**), we examined whether SRSF7, SRSF3, and SRSF5 formed sub-micron scale clusters. MRPS measurements showed that SRSF7, SRSF3, and SRSF5 formed clusters that were ∼275 – 300 nm in diameter (**Figure 4J**). These sizes remained essentially invariant across a wide range of protein concentrations, although the abundance increased with increasing concentration (**Figure S4G**). Confocal imaging using SRSF3, SRSF7, and SRSF5 labeled with AlexaFluor 405 at the N-terminus showed the formation of sub-micron scale clusters for each of the proteins (**Figure 4K**). These observations indicate that SALR-like interactions drive the clustering of microphases of SRSF7, SRSF3, and SRSF5.

### Microphase separation and clustering are modulated in mixtures of SRSFs

Since a mixture of different SRSFs can be present in nuclear speckles, we asked if mixtures of SRSF1 and one of SRSF7, SRSF3, or SRSF5 could affect SRSF1 microphase separation and the clustering of microphases in vitro. To calibrate expectations for mixtures of different pairs of SRSFs, we computed the normalized interaction parameters Δ_XY_ for pairs of domains from SRSF1 and SRSF3 (**Figure 5A**). The inter-RRM interactions are nearly ideal; the RRM-IDR interactions are equivalently attractive, whereas the inter-IDR interactions are differently, albeit strongly repulsive.

**Figure 5.**
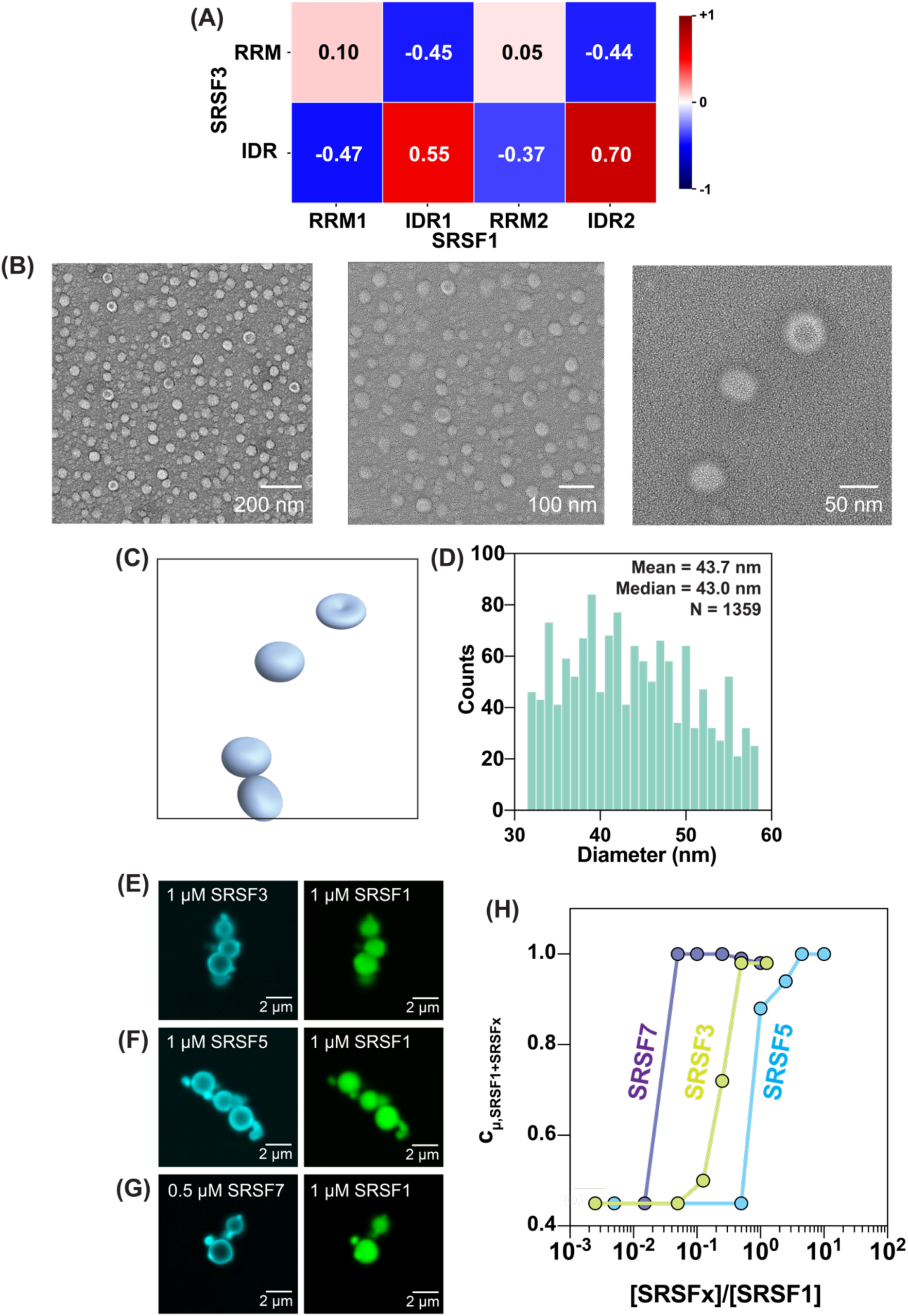
Hierarchies of interactions and structures characterize mixtures of SRSFs. **(A)** Computed interaction matrices for pairs of domains drawn from SRSF1 and SRSF3. **(B)** QF-DEEM images at different magnifications of nanoscale structures of microphases formed in mixtures of SRSF1 with SRSF3. **(C)** Three-dimensional rendering of the morphologies observed in panel **(B)**. The rendered images show the types of biconcave (same as **2D**) and spheroidal structures (same as **4H**) that were observed. **(D)** Histogram of diameters (in nm) constructed from the analysis of 1359 distinct particle images. **(E)** Confocal images of sub-micron scale structures formed in mixtures of SRSF3 and SRSF1. **(F)** Confocal images of sub-micron scale structures formed in mixtures of SRSF5 and SRSF1. **(G)** Confocal images of sub-micron scale structures formed in mixtures of SRSF7 and SRSF1. In panels (E), (F), and (G), the concentrations of SRSF3, SRSF5, and SRSF7 were below their c_µ_ whereas the concentration of SRSF1 was above its c_µ_. **(H)** Impact of each of SRSF3, SRSF5, and SRSF7 on the c_µ_ of SRSF1.

The computed interaction patterns suggest that in mixtures of SRSF1 and SRSF3 the attractive interactions involve RRMs and the IDRs, whereas the repulsions are mainly among the IDRs, akin to the individual interaction patterns of SRSF1 (**Figure 1G**) and SRSF3 (**Figure 4B**). The attractive RRM-IDR interactions are strongest for SRSF1 and weakest for SRSF3, being weaker than the RRM-IDR interactions among the two proteins (**Figure 1G, 4B, 5A**). The inter-IDR repulsions are strongest among the domains of the two proteins (**Figure 5A**). These patterns suggest an organization that maximizes attractions among SRSF1 as well as between SRSF1 and SRSF3 while minimizing inter-IDR repulsions. Thus, for a mixture of SRSF1 and SRSF3, this should result in strong SRSF1 interactions at the center to form microphases and SRSF3 preferentially accumulating on the surfaces of SRSF1 microphases, while engaging in attractive IDR-RRM interactions.

We used QF-DEEM to analyze the nanoscale structures formed in 2:1 molar mixtures of SRSF1 and SRSF3. The concentrations were chosen so that 2 µM SRSF1 is above its c_µ_ whereas 1 µM SRSF3 is below its c_µ_ (**Figure 4E**). In the QF-DEEM images, we observed two distinct morphologies, namely, biconcave discs and spheroids (**Figure 5B, 5C**). The mean and median size were 43.7 nm and 43.0 nm, respectively (**Figure 5D**). The biconcave structures were akin to those of the SRSF1 microphases (**Figure 2C**). The spheroidal structures could be a result of SRSF3 molecules filling the indentation in biconcave structures observed for SRSF1 by accumulating on the surfaces. This proposal derives from the patterns of inter-versus intramolecular interactions summarized above.

Next, we investigated the spatial organization of SRSF molecules in binary mixtures using confocal imaging of clusters of microphases. We analyzed a mixture comprising 1 µM SRSF1, which is above the c_µ_ of SRSF1, with other SRSFs at concentrations below their respective c_µ_ values. In these experiments, we used a low concentration of AlexaFluor-labeled protein and an excess of unlabeled protein to minimize any artifacts from fluorescent labeling. Across all fields-of-view, we observed preferential organization of SRSF3 on the exterior and SRSF1 on the interior of the clusters (**Figure 5E, S5A**). Similar spatial organization was observed in clusters formed in 2:1 mixtures of SRSF1 with SRSF7 (**Figure 5F, S5B**) and 1:1 mixtures of SRSF5 (**Figure 5G, S5C**), respectively. Furthermore, we observed that the size of the clusters ranged from sub-micron scale to larger than a micron in diameter, although adjacent clusters still did not coalesce.

Finally, we asked if the interactions of SRSFs with SRSF1 affected the driving forces for microphase separation of SRSF1. For this, we measured c_µ_ values in mixtures using RALS at different molar ratios of SRSFx-to-SRSF1 (x=3, 5, and 7) (**Figure 5H**). Even sub-stoichiometric amounts of the non-SRSF1 proteins were found to be sufficient at driving an increase in the c_µ_ of SRSF1, indicating that driving forces for microphase separation are weakened by interactions with other SRSFs. The c_µ_ values increased with increasing concentrations of the non-SRSF1 proteins, and they plateaued at ∼1 µM, which is approximately two times the intrinsic c_µ_ of SRSF1 (**Figure 5H**). Of the SRSFs studied here, SRSF7 is most efficient at increasing the c_µ_, followed by SRSF3 and SRSF5. These data suggest that in addition to forming their own microphases, albeit at threshold concentrations that are higher than the c_µ_ of SRSF1, the non-SRSF1 proteins weaken the driving forces for SRSF1 microphase separation, doing so in a protein-specific manner. The implication is that the stability of SRSF1 microphases can be titrated via interactions with other SRSFs.

### TDP-43, a non-SRSF speckle-associated protein, also undergoes microphase separation

TDP-43 is a speckle-associated protein, which forms TDP bodies that overlap with or co-localize with nuclear speckles under overexpression conditions ^86^. It has been reported that phase separation of TDP-43 is promoted by arginine-rich dipeptide repeat proteins (poly-GR and poly-PR), which resemble the IDR2 of SRSF1 that is enriched in Ser/Arg dipeptide repeats ^87^. TDP-43 also has been shown to co-localize specifically with sites of active splicing in mammalian neurons ^88^. Like SRSF1 and hnRNP-A1, TDP-43 has two RRMs. In addition, it has a folded N-terminal oligomerization domain (NTD) ^87,89^, and a prion-like C-terminal low complexity domain (LCD)^90^ (**Figure 6A**). The SEPs of the two RRMs (**Figure 6B, S6A**) are different from those of RRMs in SRSFs and hnRNP-A1. RRM2 of TDP-43 has a negative median SEP at-3.66 kcal/mol**·***e* (**Figure 6B, S6A**), and the median hydrophobicity of the two RRMs is higher (**Figure S6B**) than that of the RRMs from SRSF1.

**Figure 6:**
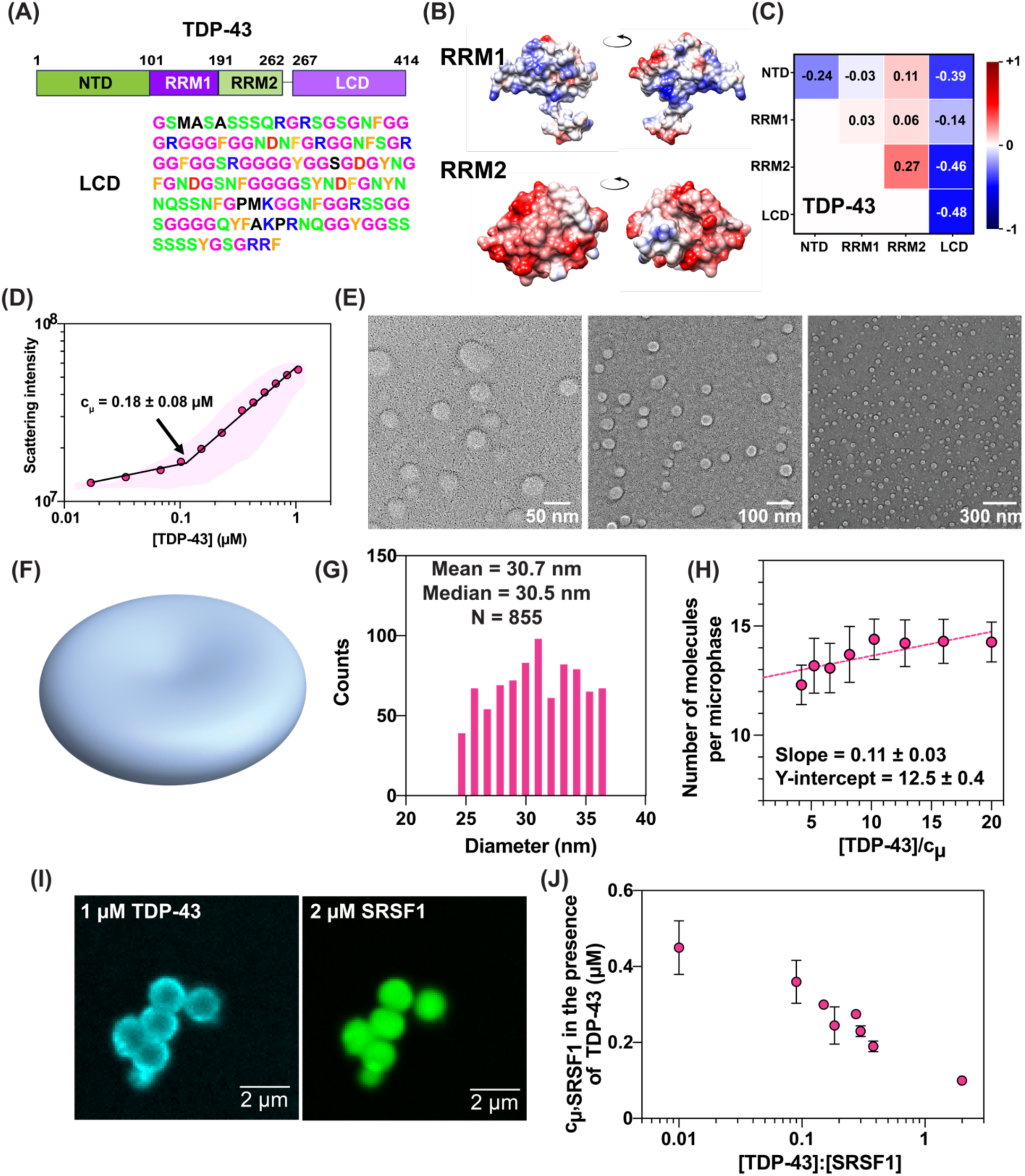
TDP-43 forms microphases and colocalizes with SRSF1. **(A)** Sequence architecture of TDP-43. Also shown are the sequence details, including the sequence of the C-terminal LCD. **(B)** Computed SEPs of each of the RRMs mapped onto the surface sites. RRM1 features patches of basic regions, and RRM2 features highly acidic faces. **(C)** Computed inter-domain interaction maps for TDP-43. **(D)** Representative RALS trace of TDP-43 showing a discontinuity at ∼0.12 µM. Results are shown from a single representative measurement (solid circles) and the spread over three technical replicates across different protein preps (pale envelope). On average, the value of c_µ_ was found to be 0.18 ± 0.08 µM. **(E)** QF-DEEM images 0.7 µM TDP-43 at different magnifications. **(F)** A rendering of the biconcave structures seen in QF-DEEM images (see **Methods**) uses a diameter of 8 units, the biconcave (dent) thickness as 1 units, and the maximum thickness of the biconcave structure near the rim as 2 units. The biconcavity thickness is larger than 0.5 used for SRSF1 (see Figure 2D) to portray dents in TDP-43 structures that are less evident than SRSF1. **(G)** Histogram of diameters of TDP-43 microphases from analysis of 855 particle images. **(H)** Results from TRFQ measurements performed over a range of TDP-43 concentrations. The concentrations of microphases were fit to a line at the plateau value and the intercept yields an estimate of the number of TDP-43 molecules per microphase to be ∼13. **(I)** Mixture of 1 µM TDP-43 and 2 µM SRSF1 show co-localization of the two proteins in size-limited clusters. TDP-43 is observed to accumulate on the surface SRSF1 clusters. **(J)** Effect of TDP-43 on the c_µ_ of SRSF1, which was measured at different molar ratios of TDP-43 and SRSF1.

Intramolecular TDP-43 interactions have been predicted between domains by sectioning the sequence into smaller regions ^91^. Here, we computed the normalized, inter-domain interaction coefficients Δ_XY_ for TDP-43 using domain boundaries delineated by the presence of folded versus disordered regions (**Figure 6C**). The most prominent inter-domain attractions are the homotypic interactions involving the LCD and NTD, and heterotypic interactions between the LCD and each of the other domains (RRM1, RRM2, and NTD). All other interactions are either repulsive (RRM2-RRM2, RRM2-NTD) or nearly ideal (RRM1-RRM1, RRM1-NTD, RRM1-RRM2). Overall, while the strengths of attractions and repulsions are weaker than those of SRSF1, the pattern of repulsions competing with attractions makes TDP-43 a tetra-block system that is more like SRSF1 and SRSF5 rather than hnRNP-A1. However, the interaction patterns are fundamentally different from SRSF1 / SRSF5.

Mutations ^92^ or aberrant posttranslational modifications ^93^ in the LCD that affect the assembly of TDP-43 could also alter the inter-domain interaction patterns that are predicted to give rise to microphase separation. It is noteworthy that previous reports pointed to assemblies of TDP-43 ranging from sub-micron scale in the absence of RNA^94^ to larger micron-scale anisosomes by RNA binding-deficient constructs that underwent coarsening over time ^95^. Some reports were based on the use of macromolecular crowders ^89^, which add depletion-mediated attractions ^96^ or the addition of specific types of RNA molecules ^97^ or chaperones ^98^ that contribute heterotypic interactions. These excipients appear to drive macrophase separation over microphase separation. Based on our computations, which lead to the classification of the TDP-43 sequence as a tetra-block copolymer, we tested for the possibility of TDP-43 microphase separation.

To test for the possibility of microphase separation of TDP-43, we performed RALS measurements, which revealed a discontinuity in scattering intensity at 0.18 ± 0.08 µM (**Figure 6D**). The rate constants determined from pyrene fluorescence lifetime measurements decreased with increasing TDP-43 concentration until c_µ_, beyond which the values of the rate constants plateaued at ∼0.93 ± 0.10 ns^-1^ (**Figure S6C**). Next, we used QF-DEEM to characterize the nanoscale structures of TDP-43 assemblies. At 0.7 µM TDP-43, which is above the measured c_µ_, we observed the formation of biconcave structures (**Figure 6E**, **6F**). The mean and median diameters of the TDP-43 microphases were 30.7 nm and 30.5 nm, respectively (**Figure 6G**). The TDP-43 microphases are 20% smaller than the microphases formed by SRSF1 and they accommodate ∼13 TDP-43 molecules per microphase as inferred from TRFQ experiments (**Figure 6H**). The number of molecules does not increase significantly for TDP-43 microphases with a slope of 0.11 molecules per micromolar increase in protein concentration.

Confocal imaging, using AlexaFluor 405-labeled TDP-43, showed the formation of irregular networks with punctate spots with low fluorescence signals, suggestive of weak TDP-43 aggregation at 2 µM, which is above the c_µ_ (**Figure S6D**). In this context, it is worth noting that TDP-43 pathology in neurons is associated with mutations within the C-terminal LCD that drives the formation of different types of aggregates ^88,99,100^. Since TDP-43 has been found in nuclear speckles, we examined mixtures of TDP-43 and SRSF1 to determine the effect on microphase separation of the two proteins. At equimolar or excess molar ratio of TDP-43:SRSF1, with both molecules being above their respective c_µ_ values, TDP-43 and SRSF1 microphases colocalized into sub-micron to micron-scale clusters (**Figure 6I**). In all fields-of-view (**Figure S6E, S6F**) and examined concentrations, the spatial organization was reminiscent of other SRSFs organizing around SRSF1 clusters (**Figure 5E-5G, S5A-S5C**). This suggests that the inter-block interactions from SRSF1 are stronger, both in terms of repulsions and attractions, when compared to the inter-protein, inter-block interactions, or the inter-block interactions within TDP-43.

Finally, we quantified how the c_µ_ of SRSF1 changed as a function of the ratio of TDP-43-to-SRSF1. The inter-protein, inter-block interactions of SRSF1 with TDP-43 enhance the driving forces for SRSF1 microphase separation (**Figure 6J**). This is seen by the systematic lowering of the c_µ_ of SRSF1 as the ratio of TDP-43 to SRSF1 reaches one. The implication is that if the TDP-43-to-SRSF1 ratio is one and the concentrations of the two species are above 0.1 µM, then the mixture undergoes microphase separation. The positive synergy between TDP-43 and SRSF1 in enhancing SRSF1 microphase separation contrasts with that of other SRSFs, which increase the c_µ_ of SRSF1. Together, these findings show how the stabilities of SRSF1 microphases can be modulated up or down by varying the levels of speckle-associated, block-copolymeric proteins such as TDP-43 or the other SRSFs.

### *MALAT1* modulates the microphase separation of SRSF1 and TDP-43

*MALAT1* is a key lncRNA that regulates splicing activity within nuclear speckles ^49,50^. It is upregulated in cancer cells ^101–103^, where it accumulates in the nucleus and localizes to nuclear speckles ^20,48,49^. *MALAT1* influences RNA processing and gene expression by serving as a hub that interacts with splicing factors such as U1 snRNP ^104^ as well as transcriptionally actively chromatin ^105^. Human *MALAT1* is ∼7k nucleotides long ^48^ and undergoes further processing to produce a 6.9k nucleotide-long transcript and a tRNA-like small RNA of 61 nucleotides^106^. *MALAT1* is predicted to compose of 200 evolutionarily conserved helices, including a 3’ triple helix ^107^, pseudoknots, tetraloops, internal loops, and long-range intramolecular interactions ^108^. *MALAT1* has been shown to interact with SRSF1 and influence its speckle localization and activity ^49,50,101^. To study the effect of *MALAT1* on microphase separation of SRSF1 and TDP-43, we produced *MALAT1* using in vitro transcription (see **Methods**). We also labeled *MALAT1* with AlexaFluor 647 at the 3’ end for fluorescence microscopy and fluorescence correlation spectroscopy (FCS).

We performed FCS using AlexaFluor 647-labeled *MALAT1* and determined the hydrodynamic radius (R_h_) to be 6.8 ± 0.7 nm (**Figure S7A**). To put this in context, the reported R_h_ of *BunV L*, a single-stranded RNA of similar length, is ∼9.2 nm in solutions where the ionic strength is more than two times lower than the ionic strength used in our measurements ^109^. These comparisons suggest that the extent of structure formation in *MALAT1* is significant, as has been shown previously ^107^, and that stable structures present within *MALAT1* help drive its compaction.

Next, we performed RALS measurements for protein-*MALAT1* mixtures comprising SRSF1 and *MALAT1* as well as TDP-43 and *MALAT1*. In the presence of a 5×10^-4^-to-1 molar ratio of *MALAT1*-to-SRSF1, we observed that the c_µ_ of SRSF1 decreases by ∼2.5-fold from 0.45 ± 0.05 µM to 0.22 ± 0.03 µM (**Figure 7A** and **S7B**). This implies that *MALAT1* promotes microphase separation of SRSF1 via preferential binding to the microphases through the mechanism of polyphasic linkage ^110–112^. Estimates based on TRFQ experiments showed that the number of SRSF1 molecules per microphase decreased from ∼50 in the absence of *MALAT1* to ∼13 molecules in the presence of *MALAT1* (**Figure 7B**). Taken together with the lowering of c_µ_, the decreased number of SRSF1 molecules per microphase suggests that *MALAT1* binding releases SRSF1 molecules, thereby contributing an entropic driving force to enhance the driving forces for SRSF1 microphase separation. It also suggests that *MALAT1* binding alters the structures of microphases and the organization within microphases when compared to SRSF1 alone.

**Figure 7.**
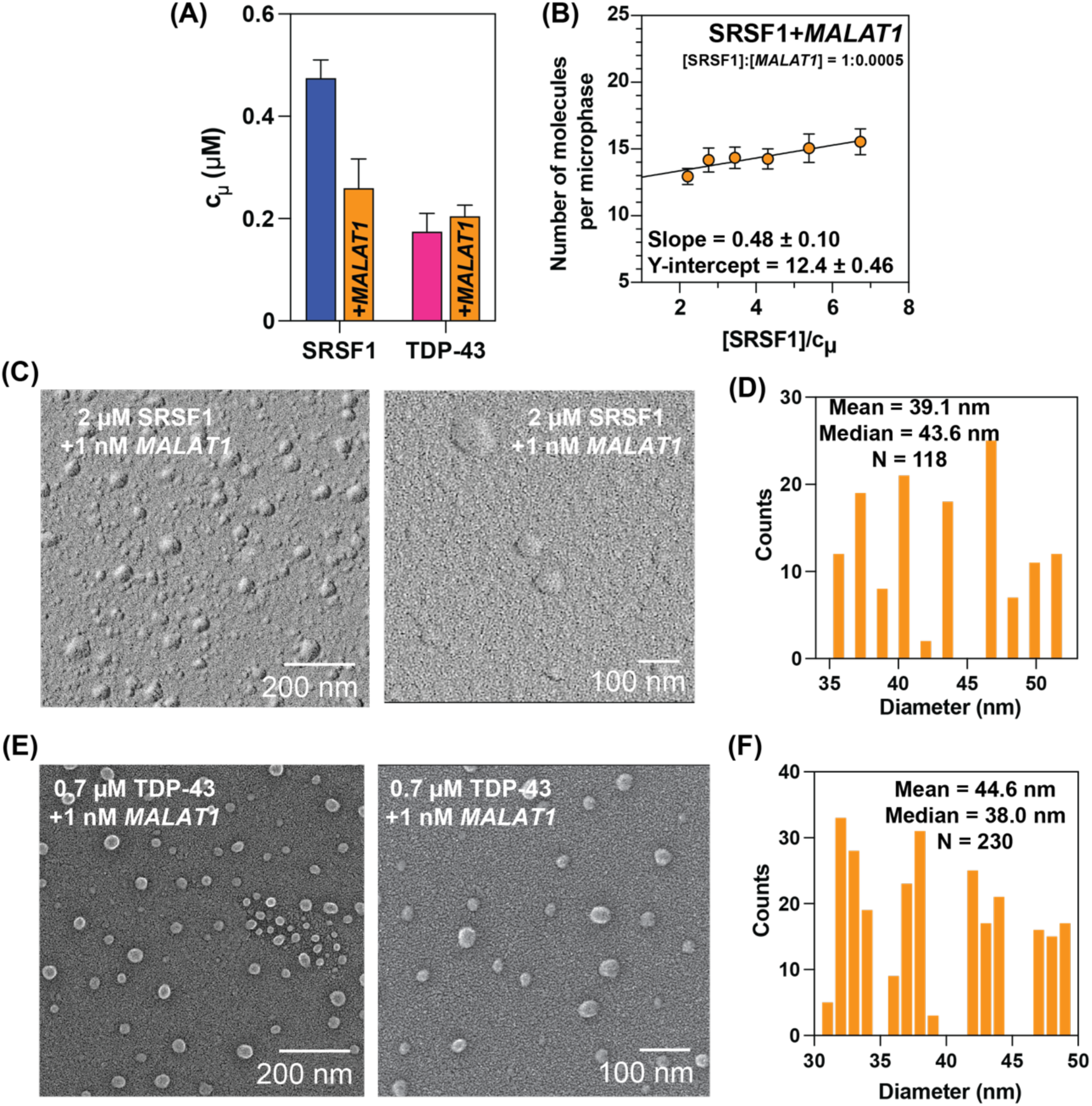
*MALAT1* binding affects microphase separation. **A.** In the presence of *MALAT1*, the c_µ_, as extracted from RALS, of SRSF1 decreased from ∼0.45 µM to ∼0.22 µM. Conversely, the c_µ_ of TDP-43 increased from ∼0.18 µM to 0.21 µM in the presence of *MALAT1*. **(B)** TRFQ experiments, performed over a range of SRSF1 concentrations at a constant molar ratio of SRSF1-to-*MALAT1* at 1-to-5×10^-4^, yielded an estimate of ∼13 SRSF1 molecules per microphase. **(C)** QF-DEEM images of SRSF1-*MALAT1* microphases show spherical morphologies, which are different from the biconcave structures observed in the absence of *MALAT1*. **(D)** Histograms collected across 118 particle images of the diameters (in nm) of the spheroids of SRSF1 microphases that form in the presence of *MALAT1*. **(E)** QF-DEEM images of TDP-43+*MALAT1* microphases. The biconcave structures formed in the absence of *MALAT1* are maintained in the presence of *MALAT1*. **(F)** Histograms collected across 230 particle images of the diameters (in nm) of the spheroids of TDP-43 microphases that form in the presence of *MALAT1*.

Using QF-DEEM, we characterized the nanoscale structures of *MALAT1*-SRSF1 microphases. In the presence of *MALAT1*, the SRSF1 microphases are spherical (**Figure 7C**) and differ from the biconcave structures observed for SRSF1 alone. The distribution of sizes is multi-modal (**Figure 7D**), and this is reflected in the different values for the mean and median, which are 39.1 nm and 43.6 nm, respectively. The distribution of sizes suggests the presence of spherical microphases of ∼37 nm, ∼40 nm, ∼44 nm, and ∼46 nm. This points to heterogeneity in the modes of binding of *MALAT1* to SRSF1 microphases.

In contrast to SRSF1, the c_µ_ of TDP-43 changed minimally (0.18 ± 0.08 µM versus 0.21 ± 0.03 µM) in the presence of the same molar ratio of *MALAT1*-to-TDP43 as *MALAT1*-to-SRSF1. This indicates that *MALAT1* binds equivalently to TDP-43 microphases and coexisting dilute phases ^110,111^ (**Figure 7A** and **S7C**). Unlike SRSF1, the biconcave structures of TDP-43 were also observed in the presence of *MALAT1* (**Figure 6E, 7E**). However, as with SRSF1, we observe a heterogeneity of microphase sizes in the presence of *MALAT1* (**Figure 7F**). The mean and median sizes are 44.6 nm and 38 nm, respectively. The dominant sizes observed were ∼32 nm, ∼38 nm, ∼42 nm, and ∼47 nm in diameter.

## Discussion

Nuclear speckles are complex membraneless bodies defined by inhomogeneous organization of multiple, compositionally specific territories that are organize around a SON-SRRM2 core ^1,7,15,19,20,24,25,50^. The physical basis for this self-organization into distinct and transient territories remains unclear ^113^. An oft-quoted hypothesis, based on specific observations, is that nuclear speckles are membraneless condensates that form via LLPS ^17,19,70,114–116^. Fei et al., reasoned that the principles of macrophase separation, adapted from models that explain multiphasic structures of nucleoli ^117^, might be relevant for explaining the speckled architectures of nuclear speckles ^20^. However, their analysis did not examine the possibility that the compositionally distinct, sub-micron scale territories could be driven by microphase separation of speckle-associated proteins such as SRSF1. Prompted by inconsistencies with an LLPS-centric mechanism, and inspired by results in the physical literature ^26^, we hypothesized that speckles comprise compositionally distinct microphases. A requisite test of this hypothesis was the demonstration that key speckle-associated proteins are block copolymeric in nature, marking them out as being different from RRM-containing proteins that are known drivers of macrophase separation ^52,53,56^.

We showed that the SRSFs studied here, as well as TDP-43, which is a different archetype of an RRM-containing speckle-associated protein, are different types of block copolymers that undergo microphase separation. The checklist for microphase separation includes the presence of a protein-specific threshold concentration c_µ_, the formation of size-limited, ordered nanoscale structures above c_µ_, and the incorporation of finite numbers (tens to hundreds) of molecules within each microphase. We discovered that the microphases can associate to form higher-order structures that either form size-limited clusters (SRSFs) or aggregates (TDP-43). Interactions that drive microphase separation are distinct from those that drive the associations of microphases. Importantly, the sub-micron scale SRSF1-rich structures observed in live cells were consistent in size with being clusters of microphases.

Unlike macrophase separation such as LLPS, where measured estimates of intra-dense-phase volume fractions suggest that macrophases concentrate >10^6^-10^7^ molecules into a single micron-scale depot ^58^, microphase separation leads to large numbers of distinct microphases, each defined by ultra-high internal concentrations, encompassing tens-to-hundreds of molecules. We can put this in context based on the parameters for the sizes of microphases and the numbers of molecules we measured within microphases. These parameters yield the following estimates: On average, there are likely to be ∼10^6^ copies of SRSF1 in a cell ^84^. Within the nucleus, this translates to a bulk concentration of 10 µM, which is above the c_µ_ values that we measured. Thus, based purely on thermodynamic considerations, microphase separation is unavoidable within the nucleus. Even if half the SRSF1 molecules are in microphases, then there are likely to be ∼50,000 SRSF1 microphases within each nucleus, with the effective SRSF1 concentration (c_eff_) in each microphase being ∼2.5 M. These findings are relevant because speckles are known to promote splicing efficiency, and a recent study suggested that even small increases in the localization of splicing factors into speckles can enable enhanced splicing activity ^118^.

The thinking is that an increased concentration of splicing factors near genes helps with increasing the splicing efficiency by driving efficient spliceosome assembly ^6,15^. Microphase separation enhances local concentrations by several orders of magnitude. Furthermore, in the presence of *MALAT1*, we discovered that the threshold concentration of SRSF1 for microphase separation is lowered, implying that *MALAT1* enhances SRSF1 microphase separation. This is achieved by preferential binding to SRSF1 that is in microphases. Interactions with *MALAT1* reduce the intra-microphase concentration of SRSF1 by ∼4-fold from 2.48 M to 0.64 M.

In accompanying work, Song et al.,^50^ tested the relevance of intrinsic and *MALAT1*-associated microphase separation of SRSFs. They showed that the phosphorylated C-terminal domain of RNA polymerase II (pCTD) binds preferentially to SRSF1 microphases, and this preferential binding is stronger than that of the unphosphorylated CTD as well as *MALAT1*. In ternary mixtures with *MALAT1* and SRSF1, pCTD was shown to displace *MALAT1* from SRSF1 microphases ^50^. These observations suggest that the recruitment of SRSF1 microphases by *MALAT1*, which stabilizes the microphases while releasing a fraction of the SRSF1 molecules from microphases, can lead to the assembly of microphases near the transcription sites of speckle-associated genes. The stronger preferential binding of the pCTD likely displaces *MALAT1* from SRSF1 microphases. The ultrahigh concentration of SRSF1 molecules within microphases, a direct consequence of their small size, is likely to be important for the regulation of splicing and the local, *MALAT1-*mediated release of splicing factors. Additionally, the high surface-to-volume ratio of microphases, which is another consequence of the small and uniform sizes, likely enables high valency, surface-mediated interactions via SRSF1 microphases.

Guo et al., showed that phosphorylation of CTD switches RNA polymerase II association from Mediator condensates to nuclear speckles. Specifically, the pCTD is incorporated into condensates formed by splicing factors such as SRSF1 and SRSF2 ^119^. These inferences, which were based on immunofluorescence microscopy, suggest that the high concentration of SRSFs in microphases serves as an attractor for pCTD ^120^. Our results provide precision regarding the local concentrations, and together with the results of Song et al.,^50^ our findings provide a physical explanation for how *MALAT1* and pCTD work together to modulate the intrinsic phase behavior of SRSF1.

We have found that the SRSF1, SRSF7, SRSF3, and SRSF5 microphases encompass ∼50, ∼46, ∼89, and ∼308 molecules, respectively. The threshold concentrations required to form these microphases are protein-specific, and microphases with fewer molecules form at lower concentrations than those with larger numbers of molecules. Tens-to-hundreds of molecules are encompassed in microphases. Thus, microphases provide a middle ground that are unlike macromolecular complexes with precise numbers of molecules or macrophases that encompass millions of molecules. The discrete number distribution of molecules within microphases is relevant given findings regarding the formation of small, microphase-sized transcriptional clusters, associated with chromatin ^121^. These clusters are short-lived, with half-lives of ∼8 s, and their small sizes make them highly sensitive to concentration fluctuations ^121^. The lifetimes of such clusters were shown to be correlated with the numbers of pre-mRNAs that are synthesized ^121,122^. As shown by Song et al., the processing of pre-mRNAs by splicing factors also utilizes high local concentrations enabled by microphase separation. To understand the contributions of finite numbers of molecules of splicing factors within microphases, we will need to find ways that toggle such systems between microphases versus macrophases.

Microphase separation has been invoked as an organizing principle for various nuclear bodies in cells. For example, the cylindrical shape of paraspeckles has been attributed to be due to micellization, a type of microphase separation, of ribonucleoprotein complexes with the lncRNA, *NEAT1* (Nuclear Enriched Abundant Transcript 1), as a scaffold. Binding of different proteins to the 5’, the middle, and the 3’ regions of *NEAT1* has been proposed to generate differential hydrophobicity along the lncRNA, thus converting protein-bound *NEAT1* into a triblock copolymer ^123^. Similarly, chromatin has been postulated to be a multi-block copolymer, where the blocks are of transcriptionally active euchromatin and inactive heterochromatin regions, promoting microphase separation ^40,42^. The binding of transcription factors such as TFEB to their target genomic loci also appear to lower the interfacial tension of condensates, thus suppressing coarsening, and driving binding-induced microphase separation ^124^. A corollary known as bridging-induced phase separation also generates size-limited DNA-cohesin clustering through DNA-cohesin-DNA bridges that give rise to microphase-like organization of chromosome-protein complexes ^125^. From a synthetic biology perspective, block copolymers made of synthetic disordered proteins such as the elastin-like (ELPs) ^45^ and resilin-like polypeptides (RLPs) ^46^ have been shown to undergo microphase separation to form distinct material properties.

Our work provides a template for the suite of methods that must be brought together to explore the possibility that nuclear proteins undergo intrinsic microphase separation that can be modulated by their RNA substrates. The accompanying work of Song et al.,^50^ provides a functional perspective on the role of microphases in speckle functions, and our work helps explain the organization of speckles into compositionally distinct territories.

### Limitations of the study

An important regulator of SRSF1-mediated interactions is the multi-site phosphorylation of Ser residues in the C-terminal IDR2 and other speckle-associated proteins ^49,62,126^. Multi-site phosphorylation can generate control over splice site selection, and the range of SRSF1 states that include unphosphorylated, hypo-phosphorylated, and hyper-phosphorylated states ^49^. Currently, we lack knowledge about how pos-translational modifications such as Ser phosphorylation impact the intrinsic and *MALAT1*-mediated microphase separation of SRSF1 and other SRSFs. Additionally, we currently lack direct probes of microphase separation in live cells that allow us to monitor how this distinctive process affects nuclear speckle functions. Despite these limitations, our work provides the first comprehensive structural studies of how SRSFs assemble. Our discovery of the relevance microphase separation, aided by multipronged biophysical approaches, helps establish a middle ground between ordered assemblies that form via site-specific binding ^127^ and multi-site cooperativity ^128,129^ on the one hand and macrophase separation on the other ^28–30,70,82,117,130^.

**Movie S1: Single-molecule tracking of slow moving SRSF1 molecules.**

Blue circles indicate localized SRSF1 molecules in each 20 ms frame, while blue lines represent single-molecule trajectories across frames. Older trajectories gradually fade over time. Raw frames containing slow dynamics (displacements less than 50 nm per 20 ms) are concatenated together. Empty frames are omitted.

**Movie S2: Single-molecule tracking of fast moving SRSF1 molecules.**

Orange circles indicate localized SRSF1 molecules in each 20 ms frame, while orange lines represent single-molecule trajectories across frames. Older trajectories gradually fade over time. Raw frames containing fast dynamics (displacements over 50 nm per 20 ms) are concatenated together. Empty frames are omitted.

## Methods

### Key resources table

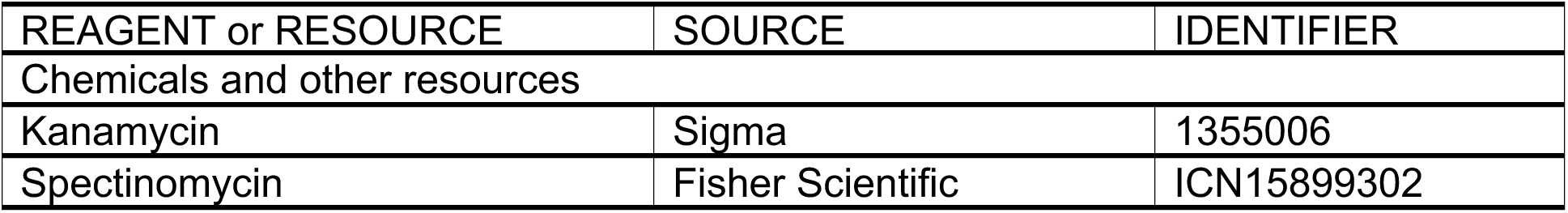
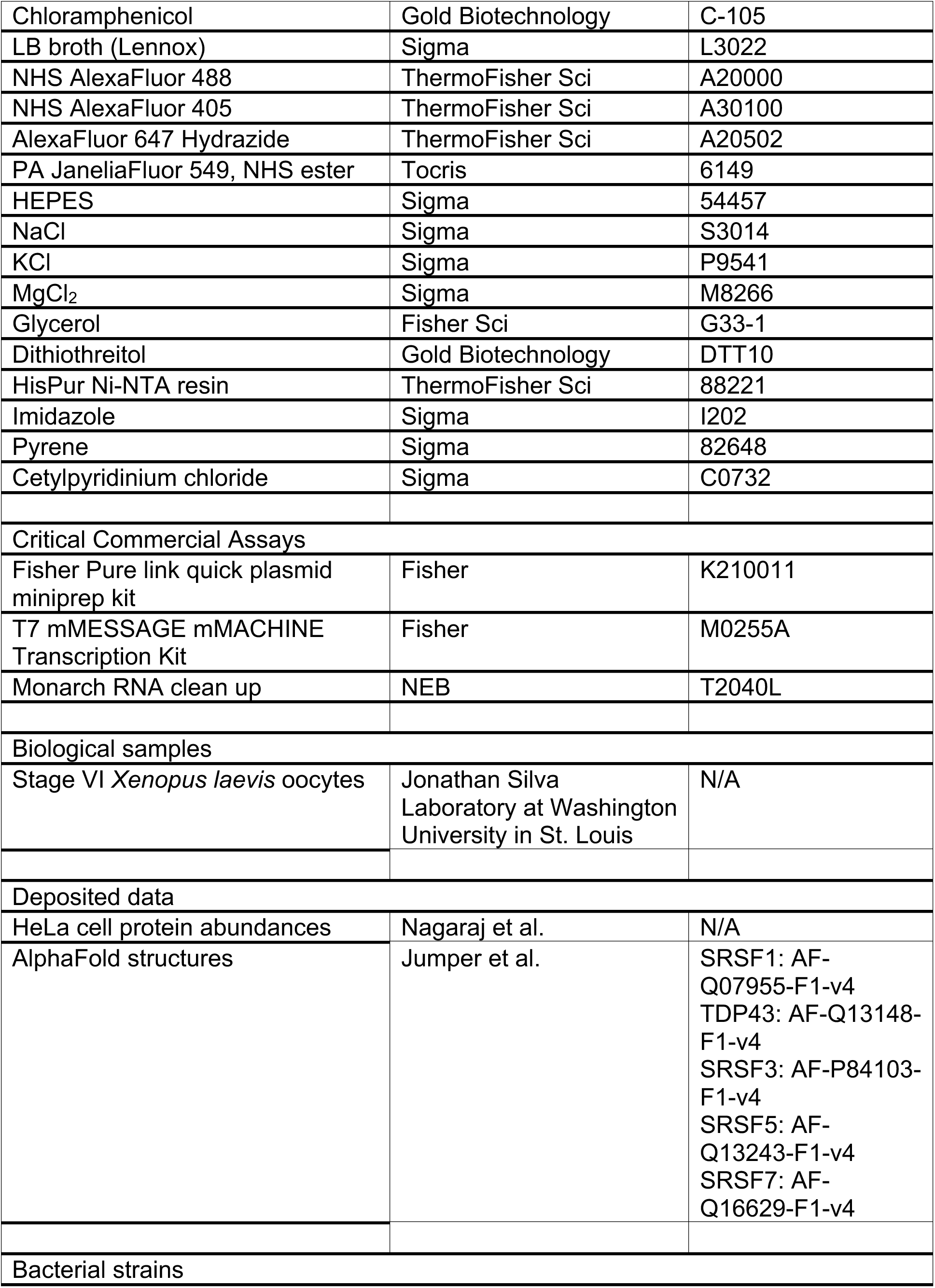
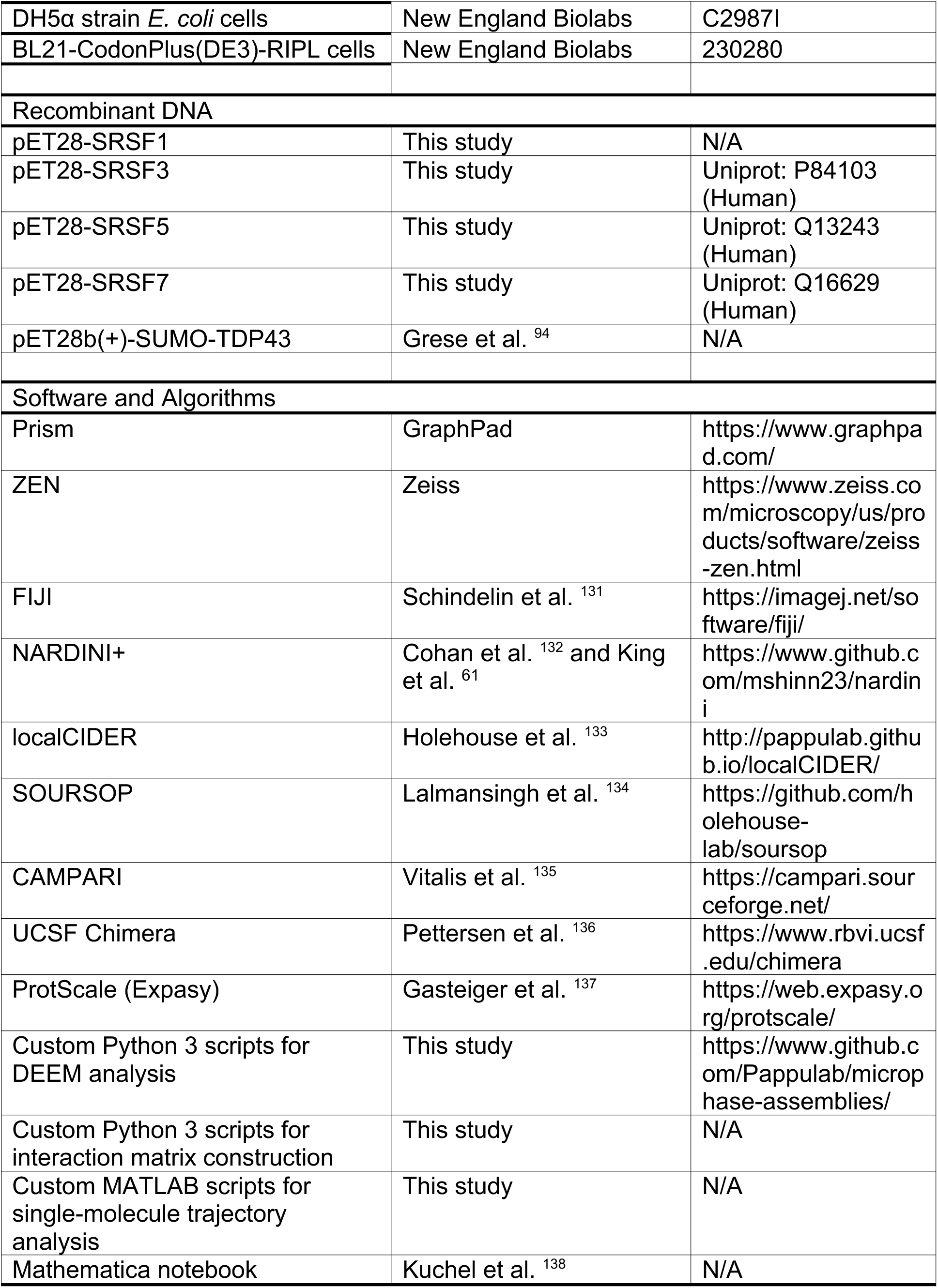

### Generation of DNA constructs

pGEMT-*MALAT1* was cloned by amplifying the *MALAT1* cDNA from pCMV-MALAT1 and ligating it into the pGEM-T easy vector (Promega, USA) according to the manufacturer’s protocol. pET28-SRSF1 was cloned by sub-cloning SRSF1 cDNA from YFP-SRSF1 into the pET28 vector by restriction digestion and ligation. pET28-SRSF3 (Uniprot: P84103), pET28-SRSF5 (Uniprot: Q13243), and pET28-SRSF7 (Uniprot: Q16629) were generated by gene synthesis (GenScript, USA). The listed pET28 constructs were transformed into the BL21-CodonPlus(DE3)-RIPL cells as described below.

Total genomic DNA was isolated using the Wizard® Genomic DNA Purification Kit (Promega, USA) as follows. Approximately 5 million HEK293T cells were harvested and washed once with PBS. Cells were lysed in 600 µL lysis buffer by vortexing followed by a 5-minute incubation with protein precipitation buffer. Lysates were centrifuged at 14,000 rpm for 4 minutes, and the supernatant was transferred to a tube with 600 µL isopropanol. After a 1-minute centrifugation at 14,000 rpm, the pellet was washed with 70% ethanol then left to dry by air for ∼15 minutes. After resuspension in 100 µL ddH_2_O, the DNA concentration and purity were measured by Nanodrop. The ratio of absorbances at 260 nm and 280 nm was measured to be ∼2.0, indicating pure DNA.

### Protein purification and labeling

#### SRSF1, SRSF3, SRSF5, and SRSF7

The His_6_-SUMO N-terminally tagged constructs were expressed in BL21-CodonPlus(DE3)-RIPL *E. coli* cells (New England Biolabs, USA) grown in LB broth (Millipore Sigma, USA), supplemented with kanamycin, spectinomycin, and chloramphenicol. Cultures were incubated at 37°C in a shaker incubator at 220 rpm until OD600 of 0.4 was reached. Cultures were then induced with 0.35 mM Isopropyl b-D-1-thiogalactopyranoside (IPTG) and incubated for additional 6-8 hours at 23°C. Cells were harvested by centrifugation at 4000 rpm for 30 minutes, washed of in Lysis Buffer (10% glycerol, 20 mM HEPES (pH 8.0), 1.25 M NaCl), and stored as pellets at-80°C.

For lysis, the cell pellet was gently resuspended in 5 mL of cold Lysis Buffer per gram of pellet with one cOmplete protease inhibitor tablet (Roche, Switzerland). 1 mg/mL of lysozyme (Roche, Switzerland) was added and incubated at 4°C for 1 hour with gentle rocking. The solution was then sonicated (Branson 550 with an L102C horn attachment) using five sets of the following 20-round cycle: 1 second on / 2 second off at 30% power. The solution was then centrifugated for 30 minutes at 10,000 rpm.

Supernatant was incubated with equilibrated 4 mL HisPur Ni-NTA resin (Thermofisher Scientific, USA) at 4°C for 1 hour. Then, the solution was poured into a manual column, and the flow-through fractions were collected. The resin was then washed with 30 mL lysis buffer and the eluted by Elution buffer (10% glycerol, 20 mM HEPES (pH 8.0), 1.25 M NaCl, 300 mM Imidazole) in 2 mL fractions. The fractions were confirmed by SDS-PAGE. Fractions containing His_6_-SUMO-SRSF1 were pooled and cleaved of SUMO tags during an overnight dialysis in the presence of 1:50 molar ratio of His_6_-tagged Ulp1 protease to His_6_-SUMO-SRSF in Cleavage Buffer (10% glycerol, 20 mM HEPES (pH 8.0), 1.25 M NaCl, 1 mM DTT (dithiothreitol)). The dialysate was incubated with 4 mL of Ni-NTA resin equilibrated with lysis buffer. Flow-through was collected, and the resin was eluted with elution buffer in 5 mL fractions. The presence of SRSF was confirmed in the flow-through and His_6_-SUMO and His_6_-Ulp1 in the elution fractions. Flow-through fractions were pooled and diluted 25-fold in dilution buffer (10% glycerol, 20 mM HEPES (pH 8.0)) to lower the total [NaCl]. This solution was further purified using a HiTrap Heparin HP 5 mL column (Cytiva, USA) on a continuous gradient with Buffer A (10% glycerol, 20 mM HEPES (pH 8.0), 50 mM NaCl), and Buffer B (10% glycerol, 20 mM HEPES (pH 8.0), 1.25 M NaCl) by FPLC (ÄKTA Pure, Cytiva, USA). Peak fractions were confirmed by SDS-PAGE, pooled, and concentrated in Amicon Ultra 3 MWCO (molecular weight cut-off) concentrator columns (Millipore Sigma, USA). Concentrated protein was aliquoted into 100 µL, flash frozen in liquid N_2_, and stored at-80°C. Concentration of protein was determined by absorbance at 280 nm. Predicted extinction coefficients^139^ for the constructs used to determine protein concentration are as follows: 24870 M^-1^cm^-1^ (SRSF1), 16960 M^-1^cm^-1^ (SRSF3), 12950 M^-1^cm^-1^ (SRSF5), 21890 M^-1^cm^-1^ (SRSF7).

#### TDP-43 (Adapted from Grese et al. ^94^)

The His_6_-SUMO N-terminally tagged construct was expressed in BL21(DE3) *E. coli* cells (New England Biolabs, USA) from the SUMO-TDP43 plasmid (pET28b(+)-SUMO-TDP43). grown in LB broth (Millipore Sigma, USA), supplemented with kanamycin. Cultures were incubated at 37°C in a shaker incubator at 220 rpm until OD600 of 0.4 was reached. Cultures were then induced with 0.35 mM Isopropyl b-D-1-thiogalactopyranoside (IPTG) and incubated for additional 6-8 hours at 23°C. Cells were harvested by centrifugation at 4000 rpm for 30 minutes, washed of in Lysis Buffer (10% glycerol, 20 mM HEPES (pH 8.0), 1.25 M NaCl), and stored as pellets at-80°C.

For lysis, the cell pellet was gently resuspended in 5 mL of cold Lysis Buffer per gram of pellet with one cOmplete protease inhibitor tablet (Roche, Switzerland). 1 mg/mL of lysozyme (Roche, Switzerland) and 10 µg/mL DNase I was added and incubated at 4°C for 1 hour with gentle rocking. The solution was then sonicated (Branson 550 with an L102C horn attachment) for 2 minutes using 30 seconds on/off cycle at 50% power. The solution was then centrifugated for 30 minutes at 10,000 rpm.

Supernatant was incubated with equilibrated 4 mL HisPur Ni-NTA resin (Thermofisher Scientific, USA) at 4°C for 1 hour. Then, the solution was poured into a manual column, and the flow-through fractions were collected. The resin was then washed with 30 mL wash buffer (10% glycerol, 20 mM HEPES (pH 8.0), 1.25 M NaCl, 50 mM Imidazole) and the eluted by Elution buffer (10% glycerol, 20 mM HEPES (pH 8.0), 1.25 M NaCl, 300 mM Imidazole) in 2 mL fractions. The fractions were confirmed by SDS-PAGE. Fractions containing His_6_-SUMO-TDP-43 were pooled and cleaved of His_6_-SUMO tags during an overnight dialysis in the presence of 1:50 molar ratio of His_6_-tagged Ulp1 protease to His_6_-SUMO-TDP-43 in Cleavage Buffer (10% glycerol, 20 mM HEPES (pH 8.0), 1.25 M NaCl, 1 mM DTT (dithiothreitol)). The dialysate was incubated with 4 mL of Ni-NTA resin equilibrated with lysis buffer. Flow-through was collected, and the resin was eluted with elution buffer in 5 mL fractions. The presence of TDP-43 was confirmed in the flow-through and His_6_-SUMO and His_6_-Ulp1 in the elution fractions. Flow-through fractions were pooled and diluted 25-fold in dilution buffer (10% glycerol, 20 mM HEPES (pH 8.0)) to lower the total [NaCl]. This solution was further purified using a HiTrap Heparin HP 5 mL column (Cytiva, USA) on a continuous gradient with Buffer A (10% glycerol, 20 mM HEPES (pH 8.0), 50 mM NaCl), and Buffer B (10% glycerol, 20 mM HEPES (pH 8.0), 1.25 M NaCl) by FPLC (ÄKTA Pure, Cytiva, USA). Peak fractions were confirmed by SDS-PAGE, pooled, and concentrated in Amicon Ultra 3 MWCO (molecular weight cut-off) concentrator columns (Millipore Sigma, USA). Concentrated protein was aliquoted into 100 µL, flash frozen in liquid N_2_, and stored at-80°C. Concentration of protein was determined by absorbance at 280 nm. Predicted extinction coefficients^139^ for the constructs used to determine protein concentration are as follows: TDP-43 is 44920 _M_-1_cm_-1.

### Labeling proteins with fluorophores

For covalent conjugation of SRSF1 or other SRSFs with NHS (h-hydroxysuccinimide ester) AlexaFluor 488 or AlexaFluor 205 (Thermofisher Scientific, USA), 1 mg of SRSF1 was dialyzed into labeling Buffer (10% glycerol, 20 mM HEPES (pH 8.3), 1.25 M NaCl) using Pierce Microdialysis Plates (ThermoFisher Scientific, USA). The dialysate was combined with NHS-Alexa Fluor dye (maintained in DMSO) at a dye:protein molar ratio of 4:1. This mixture was incubated under gentle rocking for one hour at 4°C in the absence of light. The mixture was then dialyzed extensively into Buffer H (10% (v/v) glycerol, 20 mM HEPES (pH 7.4), 50 mM KCl, 5 mM MgCl_2_) to remove unincorporated dye. Concentrations of protein and AlexaFluor 488 were determined by absorbance measurements (ε280 = 27850 M^-1^cm^-1^, ε495 = 73,000 M^-1^cm^-1^). Labeling efficiency was determined as the molar ratio of dye to protein and ranged between 0.4 –0.6. Labeled SRSF1 was aliquoted into single-use volumes (2 µL), flash frozen in liquid N_2_, and stored at-80 C.

### In vitro transcription and 3’ RNA labelling

For all RNA reagents, plasmids containing the desired RNA transcript were linearized via incubation with 5% v/v restriction enzymes (*MALAT1* – SalI) and 10% v/v CutSmart Buffer (New England Biolabs, USA) at 37°C for 4 hours. Linearization was confirmed by gel electrophoresis of DNA samples in a 1% agarose gel with non-digested DNA as controls. Linearized plasmids were purified via PCR clean-up kit (IBI Scientific, USA) per manufacturer manual. In vitro transcription was performed for each transcript using the mMESSAGE mMACHINE Transcription Kit (ThermoFisher, USA) with either SP6 or T7 polymerase as indicated. Transcribed RNAs were purified using the Monarch RNA clean up kit (New England Biolabs, USA). The purity and molecular weight of RNA were confirmed using gel electrophoresis. Confirmed RNA transcripts were aliquoted into single use volumes (2 µL), flash frozen with liquid N_2_, and stored at-80°C. *MALAT1* transcripts were labelled with AlexaFluor 647 hydrazide as described previously ^61^.

### Right angle light scattering (RALS)

SRSF1 was dialyzed against the experimental buffer (10% glycerol, 20 mM HEPES (pH 7.4), 50 mM KCl, 5 mM MgCl_2_) using Pierce Microdialysis Plates (ThermoFisher Scientific, USA) for each set of experiments. The intensity of static right-angle light scattering at 320 nm (I) was measured at each titration point using PTI Quantamaster (Horiba Scientific, Japan). The intersection of linear fits to I versus [SRSF1] (log-log plot) identified discontinuity points in the concentration dependence of the scattering intensity, which is indicative of an assembly formation and/or phase transition.

For each experiment, we report results using at least two biological and three technical replicates.

Optimal linear fits were determined using a modified jackknife approach ^140^. In summary, each dataset was subjected to two independent series of linear fits, with one series of fits for the low concentration arm (LCA) and the second series of fits for the high concentration arm (HCA). For each arm, the fitting procedure was initiated with a linear fit to the four lowest or highest concentration data points, respectively, and the root mean square error (RMSE) of each fit was recorded. Following these initial fits, the next highest or lowest concentration data point was added to the respective LCA or HCA data set, the expanded data sets were re-fit, and the new RMSEs were recorded. This process was continued, expanding the fitted dataset by one data point at a time, until all the points in the full dataset were included in both the LCA and HCA linear fits. The intersections of each pair of LCA and HCA best fits were recorded, and the average intersection point for all best fits, for all trials at a given protein concentration, were determined. This is the value reported as the threshold concentration for microphase separation.

### Time-resolved Fluorescence Quenching (TRFQ)^69^

Samples were prepared with noted concentration of protein and a constant concentration of 1 µM pyrene and 0.4 µM cetylpyridinium chloride. The 340 nm LED was used as an excitation source with a 365 nm long pass filter in an EasyLife X Lifetime Fluorescence Spectrometer (Horiba Scientific, Japan).

The fluorescence decay curves were recorded using the EasyLife X software and fitted by Eq. 1 in Prism 10 software (GraphPad, USA).

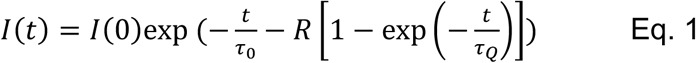

*I*(*t*) and *I*(0) are fluorescence intensities at time *t* and zero, respectively. τ_0_, τ_Q_, *R* are three parameters representing the results of the nonlinear fit. τ_0_ is pyrene lifetime in the absence of quencher. τ_Q_ is pyrene lifetime in the presence of quencher. In the presence of quencher, the fluorescence decay is described by the Eq. 1 with *k*_Q_ = 1/τ_Q_ being the rate constant for inter microphase quenching. The extracted parameter *R*, whichis equal to *c*_Q_/*c*_mic_,which is the ratio of the quencher concentration *c*_Q_ to the concentration of microphases *c*_mic_ The number of molecules per microphase, *N*, can be calculated using the following relationship (Eq. 2):

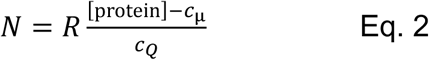

The pyrene concentration was chosen to be less than 0.05 × quencher concentration which avoids the formation of pyrene excimers. Individual solutions at a given surfactant concentration were prepared by dilution from the solution with the largest surfactant concentration. For each solution, the fluorescence decay curves were recorded and analyzed, and the plots of the number of molecules per microphase were generated for each of the protein concentrations.

### Fluorescence Confocal Microscopy

Confocal microscopy imaging performed on the Eclipse Ti2 microscope (Nikon, Japan) with a Yokogawa CSU X1 disk module and a LunF laser launch equipped with 405 nm, 488 nm, 561 nm, and 647 nm lasers, using a 60X, 1.4 NA Apo oil immersion objective (Nikon, Japan) and an Orca Flash 4.0 CMOS camera (Hamamatsu, Japan). All images were captured at room temperature using NIS-Elements software (Nikon, Japan) and saved as 16-bit ‘.nd2’ files. Images within a data set were taken with identical imaging parameters ensuring that signal was not saturated (averaging, binning, and projecting were not used). All images of purified SRSF1 and RNA constructs shown are representative crops of one or a few entities (e.g., a condensate) where the brightness and contrast have been optimized. Line scans were obtained using ImageJ and plotted using Prism 10 (GraphPad, USA). Samples were prepared with 1:250 molar ratio of labeled-to-unlabeled SRSF1 and 1:50 labeled-to-unlabeled RNA and imaged in silicone wells mounted on a coverslip.

### Single-molecule imaging of SRSF1 with PA-JF-549

We covalently labeled SRSF1 protein with photoactivatable Janelia Fluor® 549 (PA-JF-549) NHS ester as described above for AlexaFluor 488 labeling. For imaging, we mixed 1 µM of unlabeled SRSF1 with labelled SRSF1 at the molar ratio of 1:50, such that each microphase is labeled, on average, with an individual fluorophore based on the determined number of SRSF1 molecules per microphase. A reduced dilution of 1:100 was applied for larger cluster imaging to mitigate the background signal. For each sample, 15 µL of solution was placed in a silicon well (Grace Bio-Labs, USA) adhered to a cover slip.

Single-molecule microscopy was performed with a home-built wide-field epi-fluorescence microscope, equipped with an oil-immersion objective (UPLSAPO100XO, NA 1.4, Olympus, Japan), a dichroic mirror (Semrock Di01-R488/561), and an emission filter (Semrock FF01-593/46). Activation (405 nm) and excitation (561 nm) lasers shared the same optical path to illuminate the sample or activate photoactivatable fluorophore, respectively. The PA-JF-549 was activated under constant low intensity (∼50 W·cm^-2^) activation wavelength. Autofluorescence and crosstalk were not observed with 405 nm activation in the emission channel. A high peak intensity (∼4088 W·cm^-2^) of 561 nm laser was used to excite activated PA-JF-549. Single-molecule images and trajectories were acquired at 50 Hz for 20,000 frames for an average of ∼300 photons per molecule in each frame. Imaging was performed in triplicate from three independent preparations.

### Single-molecule image analysis: localization and tracking

Single-molecule blinking, and movements were observed within the acquired image stacks. Direct projections were created by summarizing all pixel information over time on finding either the maximum value (maximum projection) or the standard deviation (std projection) of the temporal intensity profile of each pixel. The maximum projection indicates the spatial distribution and emission intensity of PA-JF-549, and the std projection indicates the switching rate between the “fluorescent” and “dark” states. Both projections were used to evaluate and optimize activation/excitation control and labeled / unlabeled protein mixture ratio for observation of single molecule behaviors.

The localization and tracking of individual molecules were performed as described^141^ by applying the Robust Statistical Estimation algorithm (RoSE)^142^. All position estimates were subsequently classified and grouped into molecular displacements. This localization algorithm enables accurate position measurements of molecules on each frame, further improving the temporal resolution. Single molecule localization depends on the sequential detection and position estimation of individual molecules in each frame. A maximum likelihood estimator was used to fit a standard point spread function to the images and measure the sub-pixel lateral positions and brightness of each molecule. The localization algorithm has ∼13 nm precision in measuring lateral position of each molecule. This theoretical limit on precision is evaluated from the average of the measured fluorescence signal (300 photons) and background (9 photons).

The dim emitters were filtered out and their positions were grouped into emitter displacements by connecting estimates between two consecutive frames. Two estimates on separate frames were classified as the same emitter if: 1) they were the closest pair of estimates between the frames, and 2) their distance, d, was within the confinement radius of 200 nm. All distances were computed in Euclidean form. We classified the moving speed of each emitter by applying a threshold to the displacement, with displacements of d < 50 nm categorized as slow and those above 50 nm as fast. The 2D histograms of fast-and slow-moving emitters were generated by counting the number of fast and slow molecules originating from each binned 25-by-25 nm^2^ region on the spatial coordinates.

Excess variance of displacements across all bins within a mask were calculated as described in Eq. 3:

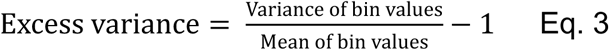

### Fluorescence Correlation Spectroscopy (FCS)

Two samples were prepared in 8-well plates (0.17 ± 0.005 mm thickness) in the experimental buffer, assumed to have the same viscosity as water: (1) a mixture of 5 nM unlabeled *MALAT1* and 0.05 nM labeled *MALAT1* and (2) 0.05 nM free dye. *MALAT1* was labeled with AlexaFluor 647 hydrazide. The free AlexaFluor 647 hydrazide was used to determine the hydrodynamic radius of the beam and used to calculate the hydrodynamic radius (R_h_) of *MALAT1* per the Stokes-Einstein Equation. The diffusion coefficient of AlexaFluor 647 hydrazide with the instrument setup used here is 3.3 x 10^-10^ m^2^s^-1^. All data were collected on a Confocor II LSM system (Carl Zeiss-Evotec, Jena, Germany) with a 40x water-immersion objective. Samples were excited at 633 nm, and emission was collected with a 650 nm cut-on long pass filter. Data for fluorescence intensity autocorrelation functions were analyzed with Zeiss Confocor II FCS software.

### Surface Electrostatic Potentials (SEPs)

The RRM structures were extracted from full-length structures of each protein from AlphaFold ^143–145^ (SRSF1: AF-Q07955-F1-v4; SRSF3: AF-P84103-F1-v4; SRSF5: AF-Q13243-F1-v4; SRSF7: AF-Q16629-F1-v4; TDP-43: AF-Q13148-F1-v4). Site-specific surface electrostatic potentials (SEPs) of RRMs were determined using “Coulombic Surface Coloring” on the solvent-excluded molecular surfaces generated by UCSF Chimera ^146^. The following default parameters were used in generating the molecular surfaces by a rolling probe sphere: probe radius of 1.4 Å to approximately represent a water molecule, vertex density of 2.0 vertices per Å^2^, and line width of 1.0 pixel in mesh surface representation. The SEP values from each vertex of the surfaces were recorded and plotted as scatterplots.

### Hydrophobicity

Hydrophobicity values were determined along the RRM sequences using a window size of 9 according to the Kyte-Doolittle scale ^147^ using ProtParam of ExPASy ^54^. The hydrophobicity values were plotted as scatterplots in Figures S1C-D, S4C, and S6B.

### Generation of biconcave disc geometry

The biconcave discs were generated as a continuous surface from a quartic equation (Eq. 4) using the Mathematica (Wolfram, USA) script from Kuchel et al^148^, while varying the diameter of the biconcave disc (d), thickness at the dent at the center (b), and maximum height at the rim (h) as noted in each figure caption. The Mathematica notebook is included in Supplementary Information. In brief, the quartic equation used was:

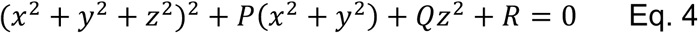

where P and Q are coefficients, R is the constant term, and x, y, and z are coordinates. Here, P, Q, and R, are defined as follows:

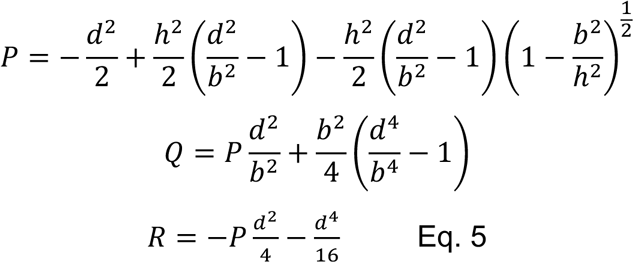

### Microfluidics Resistive Pulse Sensing

Microfluidics Resistive Pulse Sensing (MRPS) measurements were performed on an nCS1 instrument (Spectradyne, USA). Samples were prepared at indicated concentrations in experimental buffer (10 vol% glycerol, 20 mM HEPES, pH 7.4, 5 mM MgCl_2_, and 50 mM KCl). 5 μL of each sample was measured using C-2000 cartridges, with a measurement range of 250–2000 nm. At least three acquisitions (triplicates) were collected and combined for analysis, with each acquisition being collected over 10 minutes. Control experiments were performed buffer in the absence of protein, where no significant abundance of particles was detected. The particle size distributions from the triplicate experiments were averaged and the resulting curve was smoothed using the LOWESS method in Prism (GraphPad, Boston, MA, USA). The plotted values for the diameter of the clusters in **Figure 4J** were determined as the diameter at the highest particle abundance (at the peak). The curves were fitted to Lorentzian distributions to accommodate the presence of low abundances away from the center of the peak. The plotted values for the diameter of the clusters were determined as the diameter at the highest particle abundance, which is located at the peak of the Lorentzian.

### Isolation and image analysis of SRSF1-positive bodies from *Xenopus laevis* oocytes^61^

Harvested oocytes were manually disrupted from clustering, then subjected to collagenase digestion (gentle rocking) for 2 hours at 18°C. Oocytes were stored in ND96 Buffer (5 mM HEPES, 96 mM NaCl, 2 mM KCl, 1.8 mM CaCl_2_, 1 mM MgCl_2_) that was filter-sterilized and supplemented with 2.5 mM sodium pyruvate (ThermoFisher, USA) and 1X penicillin-streptomycin (MilliporeSigma, USA) at 18°C. Healthy stage VI oocytes were selected and injected using freshly pulled microneedles (Drummond, USA) and an injector (Nanoinject II, Drummond, USA). A total of 23 nL of mRNA in ddH2O (typically at a total mass of 20 ng) encoding for GFP-SRSF1 were injected into each oocyte. Injected oocytes were stored individually in wells in a 48-well polystyrene SterileTissue Culture Plates (Fisher Scientific, USA) supplemented with ND-96 buffer at 18°C for at least 18 hours to allow expression and localization of exogenous protein. Immediately prior to imaging, the germinal vesicles were manually dissected in mineral oil and mounted on a glass slide with 6 µL of mineral oil. A 22 x 22mm glass coverslip was overlaid onto the sample and then immediately imaged. This procedure was carried out for at least two separate harvests of oocytes and similar results were obtained across these biological replicates. All images shown are of a single Z-slice that is representative and is at least 2 mm above the coverslip; brightness and contrast have been optimized. For more details, see Image collection and analysis section.

The diameters of SRSF1-positive bodies were determined by using Fiji^149^. Apparent bodies smaller than 32 pixels (0.10835 mm/pixel) were omitted due to falling below resolution limits. Using the “Analyze particle” and the “Particle manager” tools, the mask for the SRSF1-positive bodies was used to obtain the area. The area was assumed to be a perfect circle and used to determine the diameter of the SRSF1-positive bodies as d = 2*sqrt(area/π). At least 100 SRSF1-bodies from at least three independent germinal vesicles were analyzed.

### Quick-freeze deep etch electron microscopy (QF-DEEM)

Samples for freeze-fracture were prepared in the experimental buffer (10 vol% glycerol, 20 mM HEPES, pH 7.4, 5 mM MgCl_2_, and 50 mM KCl). 3 µL drops of sample were added onto a glass chip supported on boiled and fixed (2% glutaraldehyde) egg white prior to rapidly pressing the sample onto a liquid helium-cooled copper block polished to a mirror finish. Frozen samples were transferred to liquid nitrogen for storage prior to replica generation.

Replicas of each sample were generated using a Leica EM ACE900 Freeze Fracture System. In brief, quick-frozen samples were transferred from liquid nitrogen to the ACE900 Freeze Fracture System. The surface was fractured at-115 to-120°C using a cold knife held at-185°C. Samples were then etched at-104°C for 3 minutes prior to shadowing with 4-5 nm Pt/C at an angle of 24°. The replica was supported with 8 nm of carbon deposited at an angle of 85° for a total of 12 nm thickness. Replicas were washed in 48 wt% HF to remove the glass support prior to washing several times with deionized water. Cleaned replica were transferred to copper grids for TEM imaging (JEOL JEM-1400 Plus) at 120 kV.

### QF-DEEM Image Analysis

Identification of microphases from.tif files were determined by the following process:

1. Particle (microphase) identification in a rectangular bounding box using a difference of Gaussians method
2. Fitting to ellipses within bounding boxes
3. Estimating the number of particles within a probe circle of a defined radius.

A Gaussian filter is applied on the image with a standard deviation of α=1 and α=3. Both operations smooth the image, which when subtracted, identify boundaries (“bounding box”) of particles. Images which have even lighting produce consistent detections while images with uneven lighting may require different intensity thresholds to be applied for sections of image. Particles within a defined range of area (e.g., 100 – 500 pixel^2^ for 10,000x magnification) are selected for further processing. The detected particles within the bounding box are fitted to an ellipse and scaled by (*ab*)^1/2^, where *a* is the semimajor and *b* is the semi-minor axis, respectively, of the fitted ellipse. Then, a circular ansatz is applied to the fitted ellipses. Particles were excluded if: 1) the fitted ellipse eccentricity was greater than 0.6; or the area of the fitted ellipse covered less than 40% of the area of the bounding box. These two criteria in combination allowed us to achieve high sensitivity in detecting areas that were circular and confidently fitted within the bounding boxes. Finally, detected particles of less than 30 nm in diameter were disregarded as unreliably resolved from the image analyses.

The average number of particles found within a probe sphere of diameter of 0.5 µm was determined using a sweep approach by generating a circular mask is generated. Partial selections of assemblies are considered as belonging to the circular mask and are accounted. The sweep is performed across the x and y ranges of the image in step size of 100 pixels. To avoid circular mask cutoffs, sweeps occur N pixels away from the image edge. A box plot of the number of particles found in all non-empty sweeps are recorded.

### Fluorescence Recovery after Photobleaching (FRAP)

FRAP was performed on the LSM 980 with Airyscan 2 (Zeiss, Germany) using a 63X high NA oil immersion objective with 488 nm excitation laser at 0.2% laser power and electron gain of 650. All images were captured at room temperature using ZEN Connect. The image was colored with Fire Lookup Table using ImageJ. The fluorescence intensity was plotted using Prism 10 (GraphPad, USA). Samples were prepared with 1:250 molar ratio of labeled-to-unlabeled AlexaFluor 488-SRSF1 and imaged in silicone wells mounted on a coverslip.

### Atomistic simulations

All-atom Metropolis Monte Carlo (MC) simulations were performed using the ABSINTH implicit solvent model and forcefield paradigm as made available in the CAMPARI simulation package (http://campari.sourceforge.net)^150,151^. The simulations utilized the abs_3.5_opls.prm parameter set in conjunction with optimized parameters for neutralizing and excess Na+ and Cl-ions ^152^. Each simulation was performed using spherical droplets with a diameter of 200 Å and explicit ions to mimic a concentration of 10 mM NaCl. Ensembles corresponding to a temperature of 340 K were used in the analysis reported in this work.

For simulations involving the folded domains, we used the predicted structures from AlphaFold (Uniprot: Q07955 (SRSF1), P84103 (SRSF3), Q13243 (SRSF5), Q16629 (SRSF7), Q13148 (TDP-43) ^153,154^). Conformations of the RRMs were initially equilibrated to make sure bond lengths and bond angles were consistent with the ABSINTH model. For simulations involving the IDRs, a single simulation was performed at 20 K to build the IDRs. Then the end PDB structure from these simulations were used to generate ten distinct full-length starting structures for ten independent simulations, each initiated using a distinct random seed. The backbone flexibility was limited by applying restraints on the starting structure, whereas the sidechain degrees of freedom were unrestrained. To free the simulations of biases from the initial coordinates, a short, high-temperature (500 K) comprising 2×10^4^ MC steps were performed. The final structures from each of the high-temperature simulations were then used as the input structures for ten independent simulations. The temperature for the production runs was set to be 340 K, which based on recent calibrations has been shown to generate conformational statistics that are congruent with various experiments ^27,155^. Each simulation comprised a total of 4.0×10^7^ MC steps that combined the full spectrum of translational, rotational, pivot, local, and concerted moves. Conformations pertaining to the first 5×10^6^ MC steps were discarded as equilibration.

The goal of the simulations was to extract pairwise inter-domain interaction coefficients, which serve as proxies for second virial coefficients. To achieve this, this each simulation comprised two domains X and Y that were either a pair of RRMs, a pair of IDRs, or an RRM and an IDR. A large spherical droplet that is over an order of magnitude larger than the dimensions of an individual domain represents an ultra-dilute system for a pair of domains. In a finite unbiased simulation, most of the inter-domain distances sampled would correspond to dissociated molecules. To overcome this problem, we applied an inter-domain distance restraint, modeled using a harmonic potential. Pairs of alpha carbon atoms located close to the centroid of each molecule were restrained to be within a shallow harmonic well, defined by a spring constant of 10 kcal/mol-Å and an equilibrium distance of 75 Å. Note that no restraints were applied on any other inter-residue or inter-atomic distances. From the ensemble of equilibrated configurations for each of the distinct pairs of domains, extracted from ten independent simulations for each pair, we extracted statistics for the inter-residue distances using MDTraj^156^ and SOURSOP ^157^. These statistics were used to compute the ensemble-averaged distances 〈*R*_XY_^(ij)^〉 between pairs of residues *i* and *j*, where the former is located on domain X and the latter on domain Y. To calibrate the observed pattern of inter-domain, inter-residue distances against a suitable prior, we performed reference simulations to model the expectations for an ideal, non-interacting pair of the domains of interest that freely samples the same simulation volume, with the presence of the weak restraint as described above. These reference simulations were performed using the inverse power potentials introduced by Tran et al ^158^. In these simulations, the non-boded interactions are such that all terms, excepting the Lennard-Jones repulsions were switched off. The exponent for the repulsive exponent was set to six, which corresponds to Maxwellian particles, and the ε was set to 0.0001. This choice was based on calibrations, which showed for IDRs, irrespective of the sequence, an exponent of six and ε of 0.0001, with all other non-bonded interactions switched off, the ensemble-averaged intramolecular, internal distance profiles showed fractal behavior whereby the distances between pairs of residues *i* and *j* 〈R*_ij_*〉 ∼ |j-i|^0.5^. This is the scaling expected for a Flory random coil. Inter-domain, inter-residue distances extracted from simulations using pairs of domains based on the inverse-power potential with an exponent of six and ε of 0.0001 were used to extract ensemble-averaged distances denoted as 〈*R*_XY,ideal_^(ij)^〉.

### Generation of computed interaction matrices

For each of domains, X and Y, the computed sets of ensemble-averaged, inter-domain, inter-residue distances denoted as 〈*R*_XY_^(ij)^〉 and 〈*R*_XY,ideal_^(GH)^〉, were used to compute excess interaction coefficients as follows:

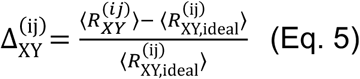

The parameter defined in Eq. 5 varies between-1 and +1. Next, we used the excess interaction coefficient for each pair of residues across the domains X and Y, as defined in Eq. 5, to compute the excess interaction coefficient Δ_XY_ using:

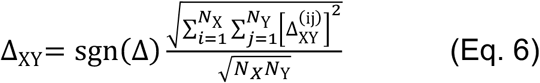

Here, Δ is the grand sum (defined as the sum over all the elements) of the upper or lower triangular matrix, where each element of the matrix is Δ_XY_^(ij)^as defined in Eq. 6. The signum or sign function sgn is such that, the sgn(Δ) is set to-1 if Δ < 0, +1 if Δ > 0, or 0 if Δ = 0 with numerical precision. The parameters *N*_X_ and *N*_Y_ refer to the numbers of residues within domains X and Y, respectively. This procedure yields values for Δ_XY_ that are normalized and signed such that-1 ≤ Δ_XY_ ≤ +1.

### Sequence feature comparison by NARDINI+

The current version of NARDINI+ ^60,61^ uses the protein sequences from the human proteome from Swissprot Homo sapiens database (May 2015, 20882 entries) ^159^. All IDR sequences within the SRSFs and TDP-43 were analyzed in terms of 90 sequence features that quantify to compositional biases and the binary patterning of pairs of residue types with respect to one another ^60,61,160^.

Non-random binary patterns were extracted using the NARDINI algorithm ^59^. This helps quantify if pairs of residue types are non-randomly distributed along the linear sequence. The non-randomness is quantified in terms of z-scores, whereby deviations from z-scores of zero quantify the magnitude and type of non-randomness. A positive z-score implies non-random segregation of the pairs of residue types in question, whereas a negative z-score implies non-random mixing of the pairs of residue types. For this analysis, we followed the approach of Cohan et al., ^59^ and grouped residues as follows: **pol** (S, T, N, Q, C, H), **hyd** (I, L, M, V), **pos** (K, R), **neg** (E, D), **aro** (F, W, Y), **ala** (A), **pro** (P), and **gly** (G).

Compositional biases were determined by analyzing 54 parameters, that include the fractions of each amino acid type (20 features); the fractions of positive, negative, polar, aliphatic, and aromatic residues (5 features); the ratios of Arg to Lys and Glu to Asp (2 features); the fraction of residues promoting chain expansion (FCE), the fraction of charged residues (FCR), the net charge per residue (NCPR), the fraction of disorder promoting residues, the hydrophobicity, the isoelectric point, and the polyproline-II propensity (7 features). Most compositional features were quantified using localCIDER^161^. The “RG Patch” feature is defined as the fraction of the sequence that is made up of patches of an Arginine-Glycine (RG) pair. Here, a patch had to have at least four occurrences of the given residue or two occurrences of RG and could not extend past two interruptions. For the various compositional features, z-scores for each IDR were calculated using the mean and standard deviation of the human IDRome. Z-scores greater than zero indicate an enrichment of the compositional feature compared to the human IDRome, and z-scores less than zero indicate a depletion of the compositional feature compared to the human IDRome.

## Acknowledgments

We thank Matthew King for help and advice on cloning of SRSF1, Harshani Pathirana (Ayala lab) for advice on purification of TDP-43, Jared Lalmansingh for help with developing methods for image analysis, Yiyang Chen for discussions on single-molecule experiments, Michael Vahey for the use of the spinning-disk confocal microscope, and Gaurav Chauhan, Souradeep Ghosh and Ruoyao Zhang for discussions on the physics of microphase separation. This project was supported by funds from the National Institutes of Health through grants K99GM152778 to MKS, R35GM124858 to MDL, T32EB019944 to YJS, and R01GM132458 the Cancer center at Illinois seed grant, ARPA-H, NSF center for quantitative cell biology to KVP, the St. Jude Children’s Research Hospital through the research collaborative on the biology and biophysics of RNP granules (to RVP), the US Air Force Office of Scientific Research (grant FA9550-20-1-0241 to RVP), and the US National Science Foundation (grant MCB-2227268 to RVP). Freeze-fracture deep-etch EM images were obtained using a JEOL JEM-1400 120kV TEM housed in the Washington University Center for Cellular Imaging (WUCCI), which is supported by Washington University School of Medicine, the Children’s Discovery Institute of Washington University and St. Louis Children’s Hospital (CDI-CORE-2015-505 and CDI-CORE-2019-813) and the Foundation for Barnes-Jewish Hospital (3770 and 4642).

## Author Contributions

MKS and RVP conceived of and designed the project following discussions about splicing factors with KVP and YJS. MKS performed all the in vitro experiments. DTT and MKS prepared samples for the QF-DEEM experiments, and DTT and GWS collected the QF-DEEM data with technical assistance and guidance from GWS. AP and MKS expressed and purified protein and RNA constructs and performed the experiments in *X. leavis* GVs. YQ and MKS performed the single-molecule experiments with guidance from MDL, who designed the experiments. VL designed and performed the all-atom simulations with inputs from MKS and RVP, under the supervision of KMR. KMR performed the NARDINI+ analysis. MKS and RVP wrote the manuscript. DTT, VL, KMR, YQ, YJS, YA, MDL, and KVP edited the manuscript. All authors read and approved of the final version.

## Supplementary Figures

**Figure S1.**
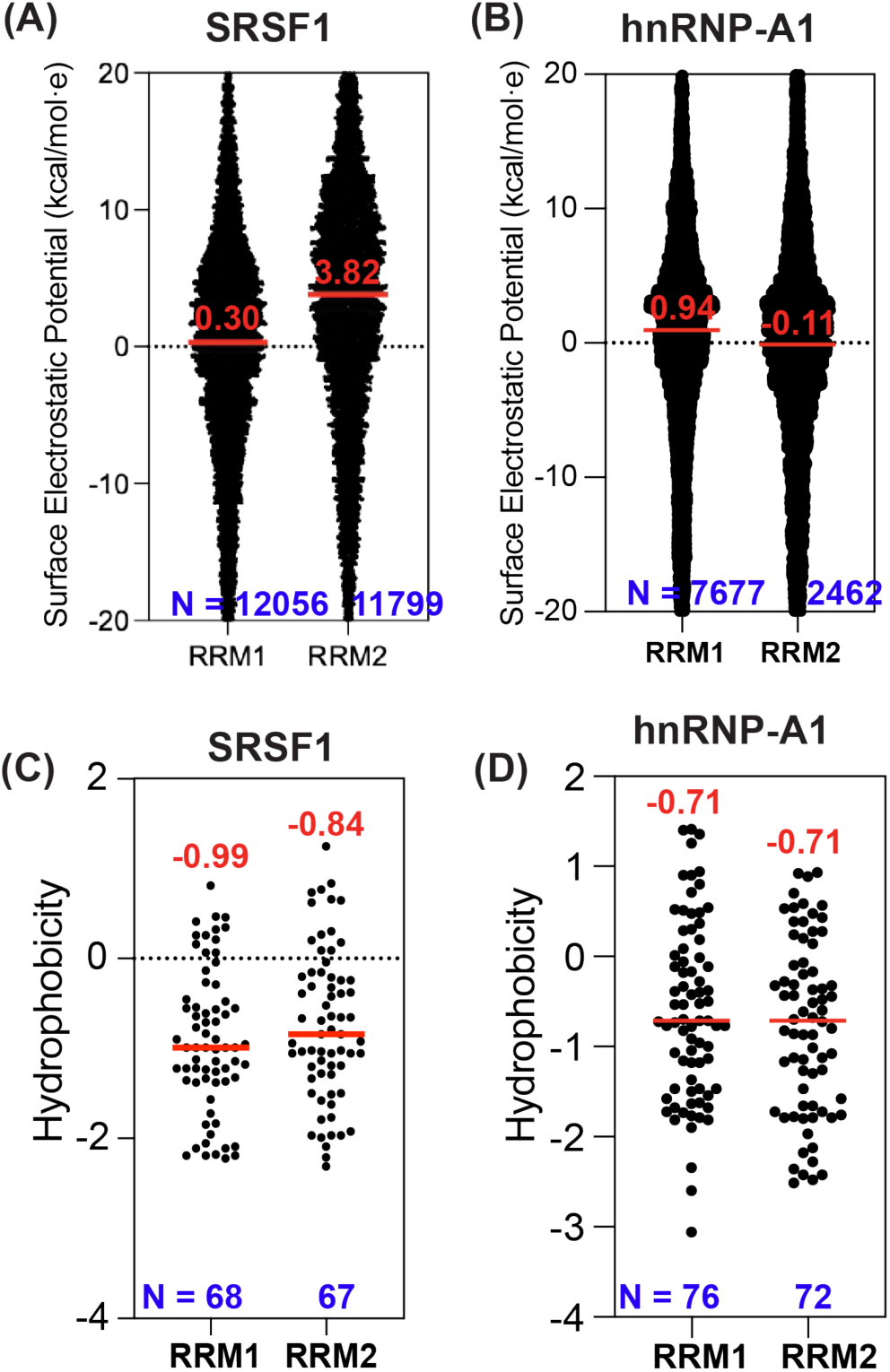
(A) Scatter plot of site-specific surface electrostatic potentials computed for the RRM1 and RRM2 of SRSF1 focused on the range between-20 and +20 to show median values. For RRM1, the median SEP is 0.30 and the potentials vary from-42.4 to +181.1 kcal/mol**·***e*. For RRM2, the median SEP is +3.82 kcal/mol**·***e* and the potential ranges from-49.9 to +181.5 kcal/mol**·***e*. The number of points for each construct is noted in blue at the bottom of the plot. **(B)** Scatter plot of site-specific SEPs computed for the RRM1 and RRM2 of hnRNP-A1. For RRM1, the median SEP is +0.94 kcal/mol**·***e*, and the potentials vary from-31.8 to +36.3 kcal/mol**·***e*. For RRM2, the median SEP is-0.11 and the potential ranges from-37.1 to +29.7 kcal/mol**·***e*. **(C)** Scatter plot of the hydrophobicity values (computed as described in the **Methods**) for RRM1 and RRM2 of SRSF1. Negative values pertain to hydrophilic linear sequence regions, whereas positive values pertain to hydrophobic regions. The red horizontal lines in each scatter plot quantify the median hydrophobicity values. **(D)** Scatter plot of the hydrophobicity values for RRM1 and RRM2 of hnRNP-A1. The red horizontal lines in each scatter plot quantify the median hydrophobicity values.

**Figure S2.**
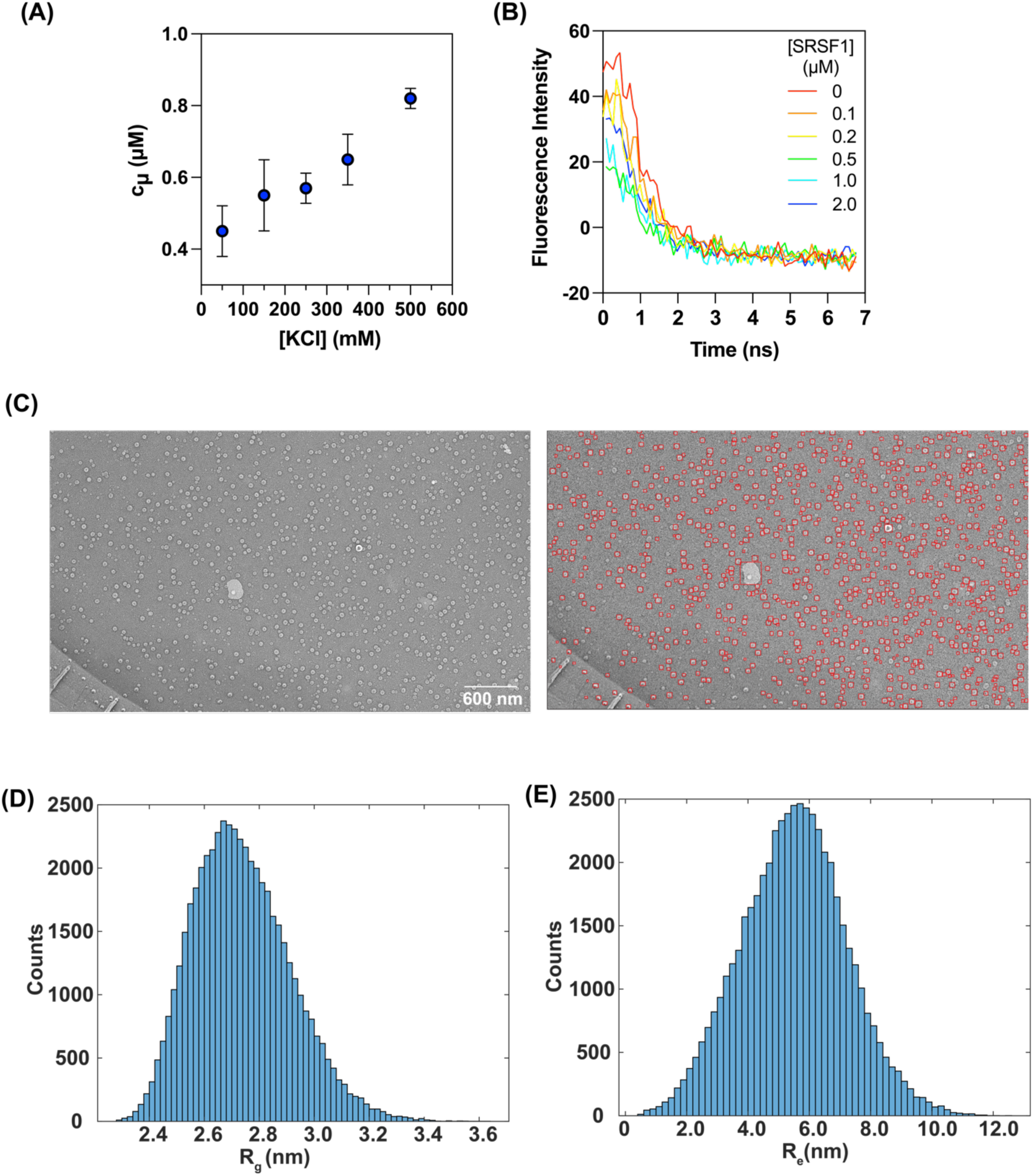
(A) The RALS extracted c_µ_ values of SRSF1 increase with increasing concentration of KCl. **(B)** Fluorescence intensity decay as a function of time measured in the presence of different concentrations of SRSF1 ranging from 0 to 2.0 µM. Pyrene is present in the solution at a constant concentration of 20 nM. **(C)** Example of QF-DEEM images that were used for quantification of sizes of SRSF1 microphases. Each particle used for the characterization of sizes is shown with a red outline (right). **(D)** Histogram of radii of gyration (R_g_) values derived from atomistic simulations (see **Methods**). The width of each bin was set to be 0.02 nm. **(E)** Histogram of end-to-end distance (R_e_) values derived from atomistic simulations (see **Methods**). The width of each bin was set to be 0.2 nm.

**Figure S3.**
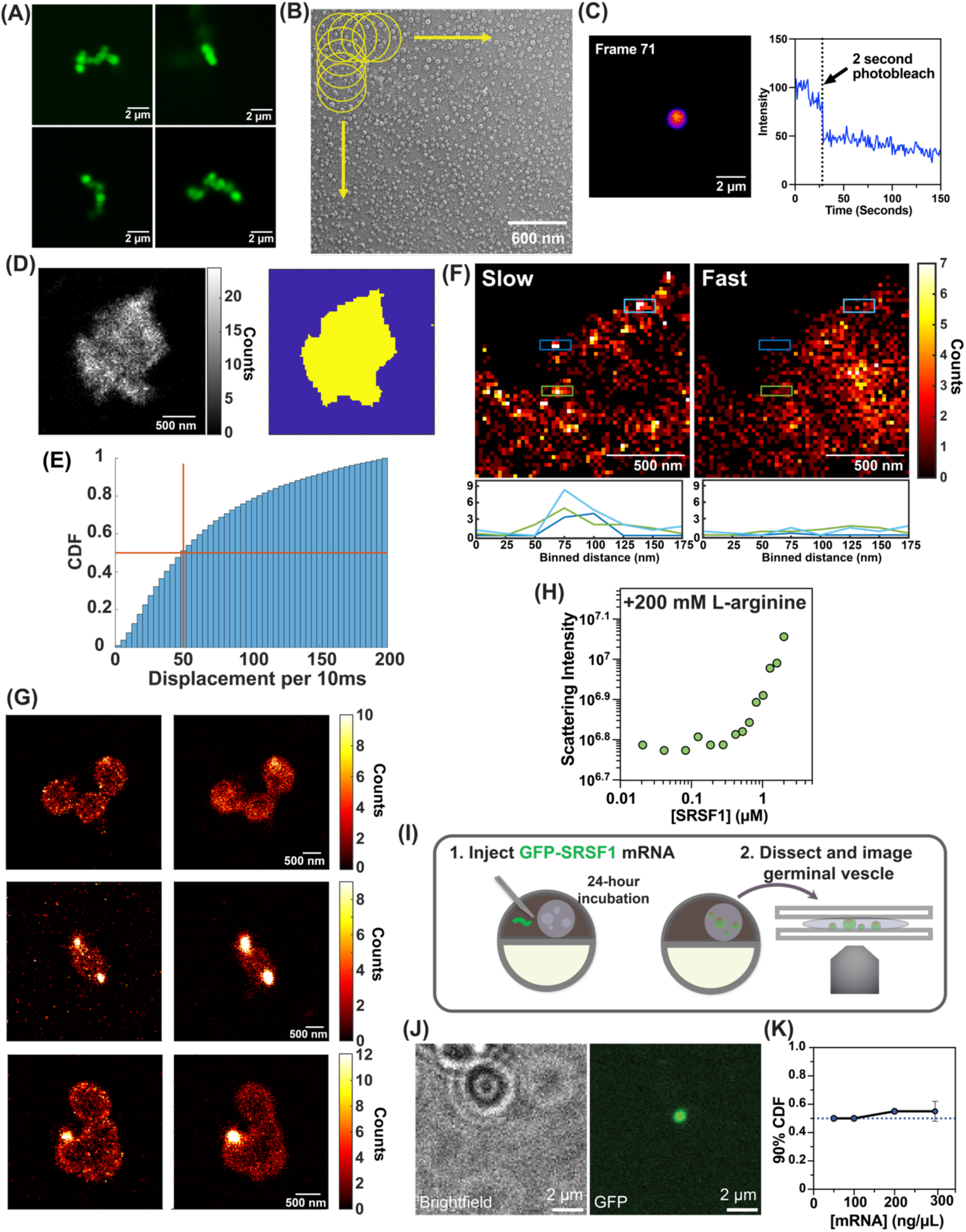
(A) Additional fields-of-view showing the confocal images of SRSF1 clusters collected at a concentration of 2 µM. **(B)** Schematic of the tiling used to estimate the number of microphases within a probe sphere with a diameter of 380 nm. **(C)** A single frame of a SRSF1 cluster from 1 µM SRSF1 sample after photobleaching (left). The fluorescence intensity was measured for 150 seconds in total. The cluster was photobleached for two seconds at the 25-second mark. Frame 71 is from the 71-second mark of the measurement corresponding to the intensity plot (right). **(D)** Single molecule image and corresponding mask of the image of a cluster comprising several localized fluorescence spots. **(E)** Cumulative distribution function (CDF) quantifying the probability of observing displacements of different magnitudes. **(F)** Histograms of displacements from slow (left) and fast (right) trajectories from insets in Figure 3F and line scan profiles below each histogram corresponding to three outlined fluorophore regions (green, light blue, dark blue). The mode of the histogram is apparent at length scales corresponding to the size of a microphase. **(G)** Histograms of fluorophores with displacements of less than 50 nm (left) or more than 50 nm (right) over each 20 ms frame for distinct sets of 2×10^4^ frames. The histograms are represented as heatmaps mapped onto the image. **(H)** RALS measurements for SRSF1 in the presence of 200 mM L-Arginine. **(I)** Schematic of microinjection of mRNA for overexpression of GFP-tagged SRSF1 in *X. laevis* oocytes. After injections, the oocytes are incubated for 24 hours, after which the germinal vesicles are extracted, placed on glass coverslips, and imaged. **(J)** Brightfield (left), fluorescence (middle), and merged (right) images of *X. laevis* oocyte overexpressing GFP-tagged SRSF1. **(K)** The diameter in µm of SRSF1-positive bodies at 90% of the CDFs (see **3L**) plotted as a function of mRNA concentration.

**Figure S4.**
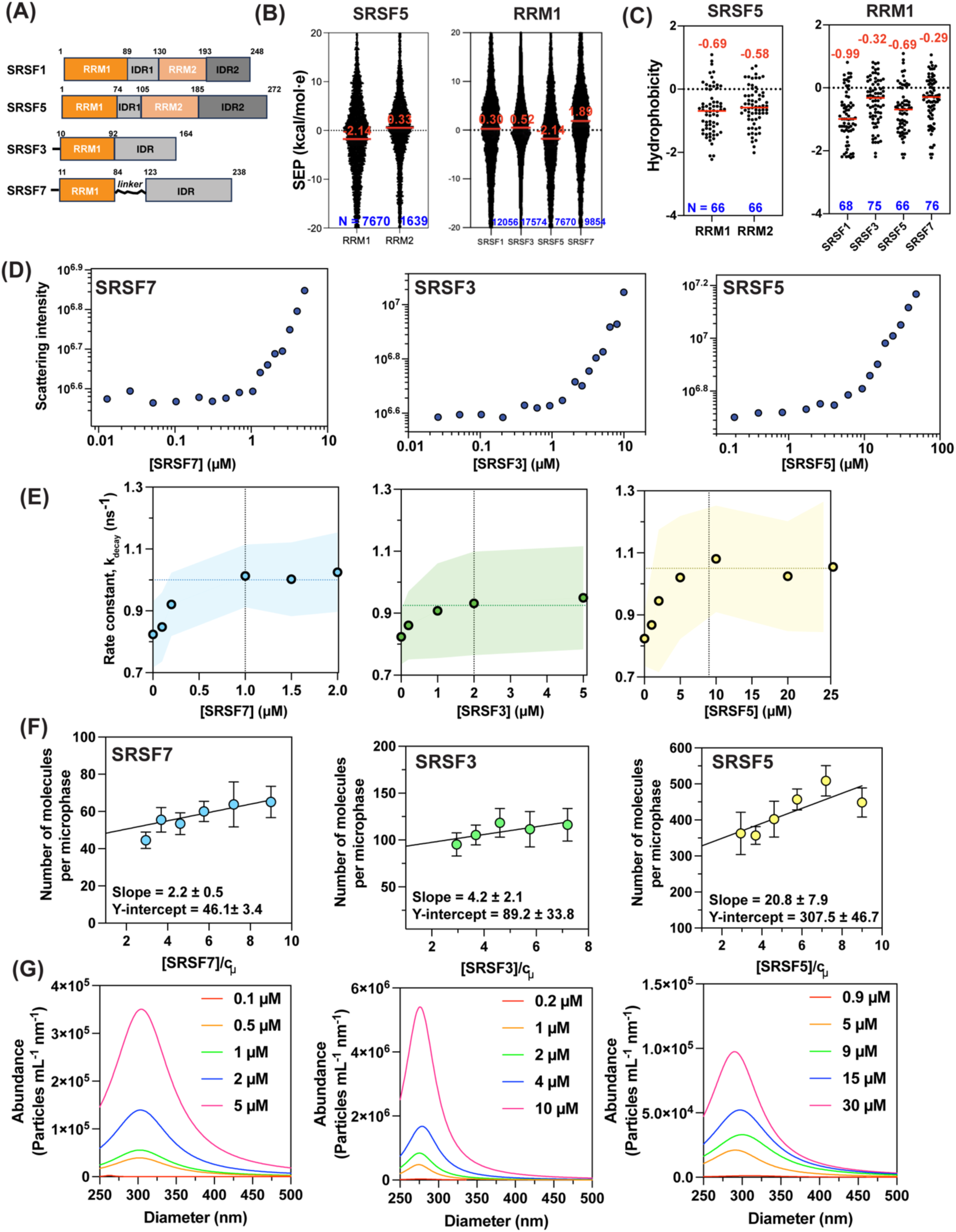
(A) Sequence architectures and domain boundaries of SRSF1, SRSF3, SRSF5, and SRSF7. **(B)** Scatter plot of SEPs with one value for each surface atom, for RRM1 and RRM2 of SRSF5 (left) and comparisons of the site-specific SEPs of RRM1 from SRSF1 and SRSF5 to the single RRM in SRSF3 and SRSF7. In each scatter plot, the red line denotes the median SEP value. **(C)** Scatter plot of computed residue-specific hydrophobicity values, with one value for each linear sequence region of nine residues, shown here for RRM1 and RRM2 of SRSF5 (left) and comparisons of the hydrophobicity values of RRM1 from SRSF1 and SRSF5 to the single RRM in SRSF3 and SRSF7. **(D)** Representative plots of concentration-dependent normalized scattering intensities measured from RALS for each of SRSF7, SRSF3, and SRSF5. **(E)** Concentration-dependent rate constants *k*_decay_ that quantify the pyrene lifetimes for each of SRSF7, SRSF3, and SRSF5. Note that the *k*_decay_ values plateau (horizontal dotted line) around the c_µ_ values (vertical dotted line) measured using RALS. Each plot shows the mean values for *k*_decay_ and an envelope that quantifies the standard error in the mean from three technical replicates. **(F)** Plot of the estimate of the number molecules per microphases determined from TRFQ assays for each of SRSF7, SRSF3, and SRSF5. The circles denote the mean of estimate of the number of molecules per microphase (see **Methods**) and the error bars are standard errors in the estimate of the mean. The intercept is used as the estimate of the number of molecules per microphase at the c_µ_. Linear regression analysis was performed as described in the **Methods**, and the slope and the Y-intercept are noted in each panel. **(G)** Size distributions extracted from MRPS data traces are shown for a range of concentrations for SRSF7, SRSF3, and SRSF5.

**Figure S5.**
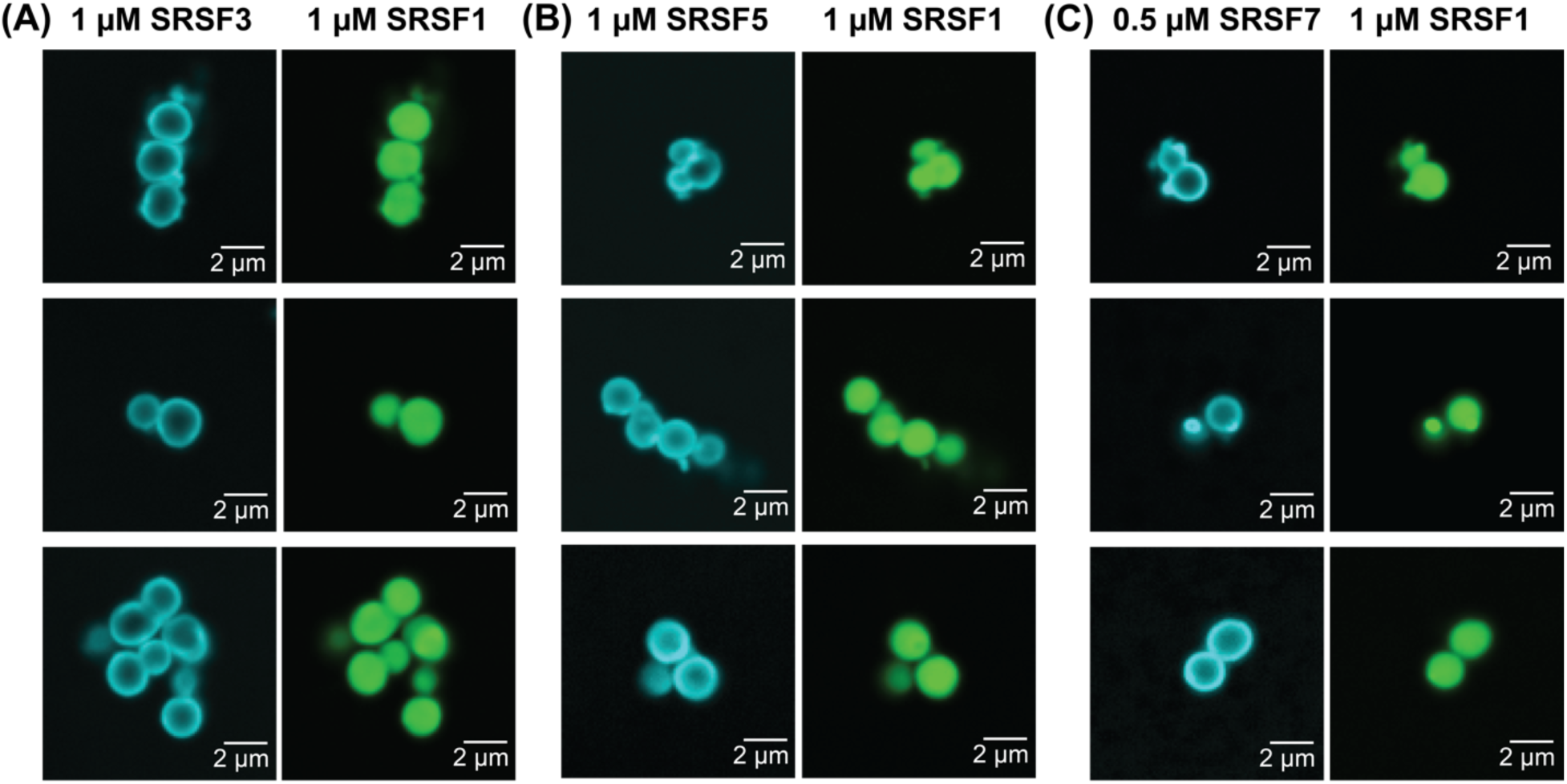
(A) Three additional fields-of-view showing the confocal images of clusters from mixtures of 1 µM SRSF1 and 1 µM SRSF3. The two panels from each row are from the same field of view for the SRSF3 (cyan, left) channel and the SRSF1 (green, right) channel. **(B)** Three additional fields-of-view showing the confocal images of clusters from mixtures of 1 µM SRSF1 and 1 µM SRSF5. The two panels from each row are from the same field of view for the SRSF5 (cyan, left) channel and the SRSF1 (green, right) channel. **(C)** Three additional fields-of-view showing the confocal images of clusters from mixtures of 1 µM SRSF1 and 0.5 µM SRSF7. The two panels from each row are from the same field of view for the SRSF7 (cyan, left) channel and the SRSF1 (green, right) channel.

**Figure S6.**
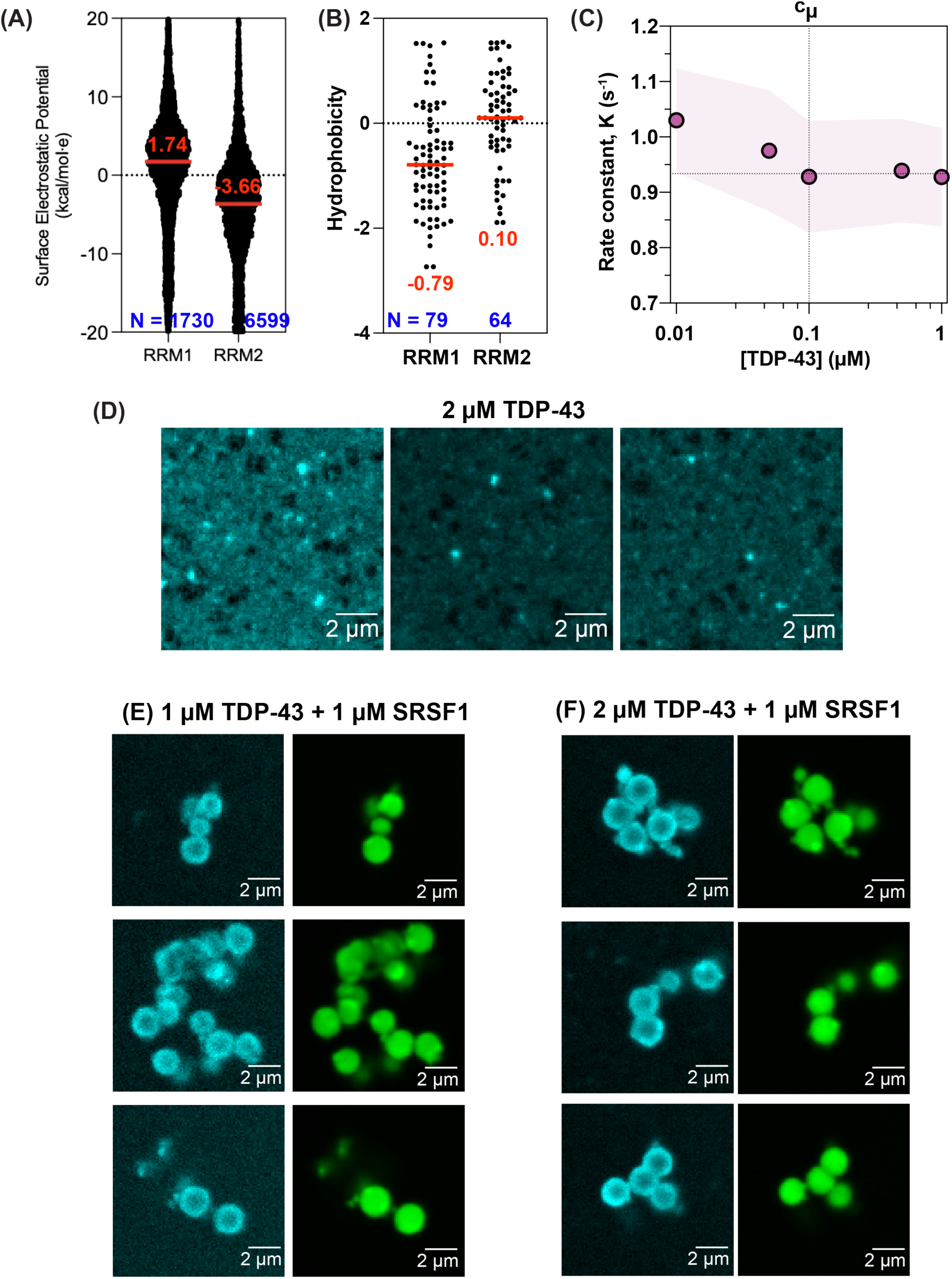
(A) Scatter plot of SEPs, with one value for each surface atom, for RRM1 and RRM2 of TDP-43 focused on the range between-20 and +20 kcal/mol**·***e* to show median values. For RRM1, the median SEP is +1.74 kcal/mol**·***e* and the potentials vary from-47.8 to +32.3 kcal/mol**·***e*. For RRM2, the median SEP is-3.66 kcal/mol**·***e* and the potential ranges from-62.7 to +29.6 kcal/mol**·***e*. The number of points for each construct is noted in blue at the bottom of the plot. In each scatter plot, the red line denotes the median SEP value. **(B)** Scatter plot of computed hydrophobicity values, with one value for each linear sequence region of nine residues, shown here for RRM1 and RRM2 of TDP-43. **(C)** Concentration-dependent rate constants *k*_decay_ that quantify the pyrene lifetimes for TDP-43. The *k*_decay_ value plateaus (horizontal dotted line) around the c_µ_ value (vertical dotted line) measured using RALS. The plot shows the mean values for *k*_decay_ and an envelope that quantifies the standard error in the mean from three technical replicates. **(D)** Confocal microscopy from different fields-of-view of irregularly shaped aggregates formed by TDP-43 at a concentration of 2 µM, which is an order of magnitude higher than the measured c_µ_. **(E)** Confocal images showing different fields-of-view of the micron scale co-localization of 1 µM TDP-43 (cyan, left) with 1 µM SRSF1 (green, right). Instead of forming the aggregates seen in panel D, we observe spatially organized micron scale clusters with TDP-43 preferentially accumulating at the surface. **(F)** Confocal images showing different fields-of-view of the micron scale co-localization of 2 µM TDP-43 (cyan, left) with 1 µM SRSF1 (green, right). Similar structures are observed to Panel E.

**Figure S7.**
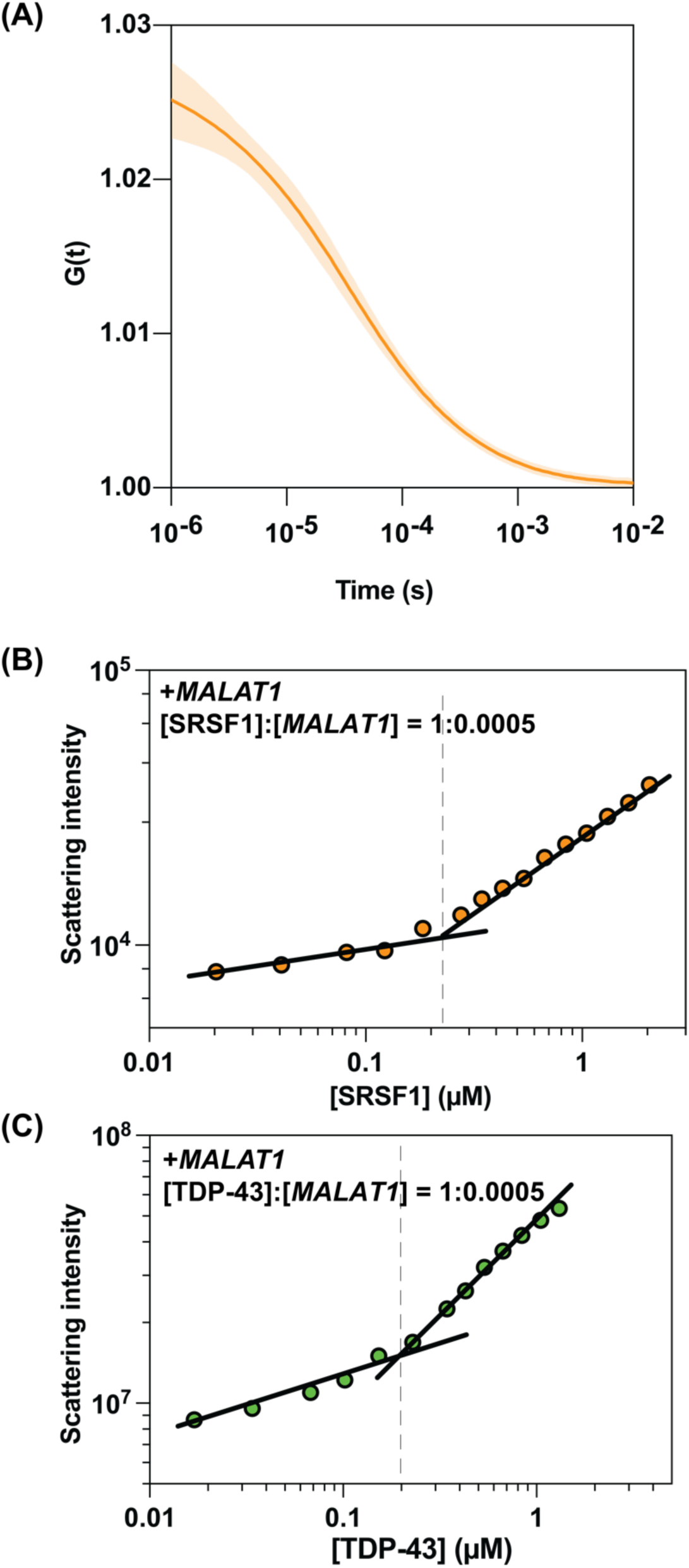
(A) Autocorrelation function from FCS measurements of AF647-labeled *MALAT1*. The mean trace and errors in the estimate of the mean from three replicates are shown as a solid curve and envelope, respectively. **(B)** Sample, concentration-dependent RALS measurement to estimate the c_µ_ of SRSF1 in the presence of a fixed, 1:5×10^-4^ ratio of SRSF1-to-*MALAT1*. The points are the measured scattered intensities and lines are drawn to join the points as guides to the eye. **(C)** Sample, concentration-dependent RALS measurement to estimate the c_µ_ of TDP-43 in the presence of a fixed, 1:5×10^-4^ ratio of TDP-43-to-*MALAT1*. The points are the measured scattered intensities and lines are drawn to join the points as guides to the eye.

